# Embeddings from deep learning transfer GO annotations beyond homology

**DOI:** 10.1101/2020.09.04.282814

**Authors:** Maria Littmann, Michael Heinzinger, Christian Dallago, Tobias Olenyi, Burkhard Rost

## Abstract

Knowing protein function is crucial to advance molecular and medical biology, yet experimental function annotations through the Gene Ontology (GO) exist for fewer than 0.5% of all known proteins. Computational methods bridge this sequence-annotation gap typically through homology-based annotation transfer by identifying sequence-similar proteins with known function or through prediction methods using evolutionary information. Here, we propose predicting GO terms through annotation transfer based on proximity of proteins in the SeqVec embedding rather than in sequence space. These embeddings originate from deep learned language models (LMs) for protein sequences (SeqVec) transferring the knowledge gained from predicting the next amino acid in 33 million protein sequences. Replicating the conditions of CAFA3, our method reaches an F_max_ of 37±2%, 50±3%, and 57±2% for BPO, MFO, and CCO, respectively. Numerically, this appears close to the top ten CAFA3 methods. When restricting the annotation transfer to proteins with <20% pairwise sequence identity to the query, performance drops (F_max_ BPO 33±2%, MFO 43±3%, CCO 53±2%); this still outperforms naïve sequence-based transfer. Preliminary results from CAFA4 appear to confirm these findings. Overall, this new concept is likely to change the annotation of proteins, in particular for proteins from smaller families or proteins with intrinsically disordered regions.

## Introduction

### GO captures cell function through hierarchical ontologies

All organisms rely on the correct functioning of their cellular workhorses, namely their proteins involved in almost all roles, ranging from molecular functions (<u>MF</u>) such as chemical catalysis of enzymes to biological processes or pathways (<u>BP</u>), e.g. realized through signal transduction. Only the perfectly orchestrated interplay between proteins allows cells to perform more complex functions, e.g., the aerobic production of energy via the citric acid cycle requires the interconnection of eight different enzymes with some of them being multi-enzyme complexes^1^. The <u>Gene Ontology (GO)</u>^2^ thrives to capture this complexity and to standardize the vocabulary used to describe protein function in a human- and machine-readable manner. GO separates different aspects of function into three hierarchies: <u>MFO</u> (Molecular Function Ontology), <u>BPO</u> (biological process ontology), and <u>CCO</u>, i.e. the cellular component(s) or subcellular localization(s) in which the protein acts.

### Computational methods bridge the sequence-annotation gap

As the experimental determination of complete GO numbers is challenging, the gap between the number of proteins with experimentally verified GO numbers and those with known sequence but unknown function (<u>sequence-annotation gap</u>) remains substantial. For instance, UniRef100^3^ (UniProt^3^ clustered at 100% percentage pairwise sequence identity, PIDE) contains roughly 220M (million) protein sequences of which fewer than 1M have annotations verified by experts (Swiss-Prot^3^ evidence codes EXP, IDA, IPI, IMP, IGI, IEP, TAS, or IC).

Computational biology has been bridging the sequence-annotation gap for decades^4-11^, based on two different concepts: (1) sequence similarity-based transfer (or *homology-based inference)* which copies the annotation from one protein to another if that is sequence similar enough because proteins of similar sequence have similar function^12^. In more formal terms: given a query Q of unknown and a template T of known function (Ft): IF PIDE(Q,T)>threshold θ, transfer annotation Ft to Q. (2) *De-novo* methods predict protein function through machine learning^5^. If applicable, the first approach tends to out-perform the second^13-16^ although it largely misses discoveries^17^. The progress of computational methods has been monitored by CAFA *(Critical Assessment of protein Function Annotation algorithms)*^9,18,19^, an international collaboration to advancing and assessing methods that bridge the sequence-annotation gap. CAFA takes place every 2-3 years with its fourth instance (CAFA4) currently being evaluated.

Here, we introduce a novel approach transferring annotations using the similarity of embeddings from language models (LMs: SeqVec^20^ and ProtBert^21^) rather than the similarity of sequence. Using embedding space proximity has helped information retrieval in natural language processing (NLP)^22-25^. By learning to predict the next amino acid given the entire previous sequence on unlabeled data (only sequences without any phenotype/label), e.g. SeqVec learned to extract features describing proteins useful as input to different tasks (transfer learning). Instead of transferring annotations from the labeled protein T with the highest percentage pairwise sequence identity (PIDE) to the query Q, we chose T as the protein with the smallest distance in embedding space (DIST^emb^) to Q. This distance also proxied the reliability of the prediction serving as threshold above which hits are considered too distant to infer annotations. Instead of picking the top hit, annotations can be inferred from the *k* closest proteins where *k* has to be optimized. In addition, we evaluate the influence of the type of LM used (SeqVec^20^ vs. ProtBert^21^). Although the LMs were never trained on GO terms, we hypothesize LM embeddings to implicitly encode information relevant for the transfer of annotations, i.e. capturing aspects of protein function because embeddings have been shown to capture rudimentary features of protein structure and function^20,21,26,27^.

## Results & Discussion

### Simple embedding-based transfer almost as good as CAFA3 top-10

First, we predicted GO terms for all 3,328 CAFA3 targets using the Gene Ontology Annotation (GOA) data set *GOA2017* (Methods), removed all entries identical to CAFA3 targets (PIDE=100%; set: *GOA2017-100)* and transferred the annotations of the closest hit (k=1; closest by Euclidean distance) in this set to the query. When applying the <u>*NK* evaluation mode</u> (no-knowledge available for query, Methods/CAFA3), the embedding-transfer reached F_max_ scores (Eqn. 3) of 37±2% for BPO (precision: P=39±2%, recall: R=36±2%, Eqns. 1/2), F1=50±3% for MFO (P=54±3%, R=47±3%), and F1=57±2% for CCO (P=61 ±3%, R=54±3%; Table 1, Fig. 1, Fig. S1). Errors were estimated through the 95% confidence intervals (± 1.96 stderr). Replacing the Euclidean by cosine distance (more standard amongst those working with embeddings, e.g. in NLP) changed little (Table 1; for simplicity, we only used Euclidean from here on). In the sense that the database with annotations to transfer (GOA2017) had been available before the CAFA3 submission deadline (February 2017), our predictions were directly comparable to CAFA3^19^. This embedding-based annotation transfer clearly outperformed the two CAFA3 baselines (Fig. 1: the simple *BLAST* for sequence-based annotation transfer, and the *Naïve* method assigning GO terms statistically based on database frequencies, here *GOA2017);* it would have performed close to the top ten CAFA3 competitors (in particular for BPO: Fig. 1) had the method competed at CAFA3.

**Fig. 1:**
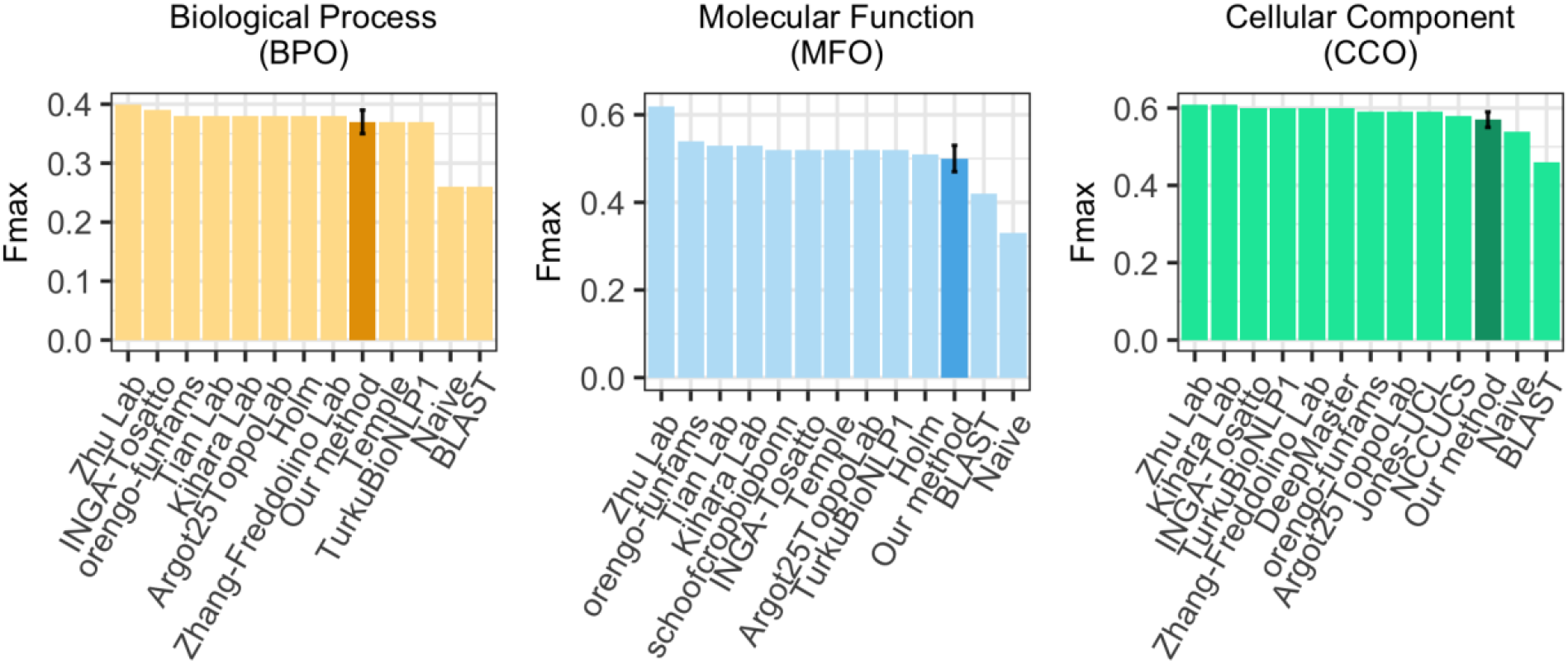
F_max_ for simplest embedding-based transfer (k=1) and CAFA3 competitors. Using the data sets and conditions from CAFA3, we compared the F_max_ of the simplest implementation of the embedding-based annotation transfer, namely the greedy (k=1) solution in which the transfer comes from exactly one closest database protein (dark bar) for the three ontologies (BPO, MFO, CCO) to the top ten methods that - in contrast to our method - did compete at CAFA3 and to two background approaches “BLAST” (homology-based inference) and “Naïve” (assignment of terms based on term frequency) (lighter bars). The result shown holds for the *NK* evaluation mode (no knowledge), i.e. only using proteins that were novel in the sense that they had no prior annotations. If we had developed our method before CAFA3, it would have almost reached the tenth place for MFO and CCO and ranked even slightly better for BPO. Error bars (for our method) marked the 95% confidence intervals.

**Table 1:** Performance for CAFA3 targets for simple GO annotation transfers^*^

| Data set | Embeddings | $F_{\max}$ | | |
| --- | --- | --- | --- | --- |
|  |  | BPO | MFO | CCO |
| GOA2017 | SeqVec | 37±2% | 50±3% | 57±2% |
|  | SeqVec (Cosine) | 37±2% | 50±3% | 58±2% |
|  | SeqVec (maximum pooling) | 35±2% | 52±3% | 58±2% |
|  | ProtBert | 36±2% | 49±3% | 59±2% |
|  | BLAST | 26% | 42% | 46% |
| GOA2017X | SeqVec | 31±2% | 51±3% | 56±2% |
| GOA2020 | SeqVec | 51±2% | 61±3% | 65±2% |
|  | ProtBert | 50±2% | 59±2% | 65±2% |
|  | BLAST | 31% | 53% | 58% |

\* Mean $F_{\max}$ values for GO term predictions using embeddings from two different language models (SeqVec or ProtBert) or sequence similarity (BLAST) for the data sets GOA2017-100 (2017), GOA2017X (2017), and GOA2020-100 (2020) used for annotation transfer (note: the notation '-100' implies that any entry in the data set with PIDE=100% to any CAFA3 protein had been removed). By default, embedding distance was assessed by Euclidean distance (Eqn. 4; exception marked *cosine*), and per-residue embeddings were pooled by global average pooling (exception marked maximum pooling). All values were compiled for picking the single top hit ( $k=1$ ) and using the CAFA3 targets from the NK and full evaluation mode<sup>19</sup>. For all simple annotation transfers (embedding- and sequence-based), performance was higher for the more recent data sets (GOA2020 vs. GOA2017). Error estimates are given as 95% confidence intervals. $F_{\max}$ values were computed using the CAFA3 tool<sup>18,19</sup>.

Performance did not change when replacing global average with maximum pooling (Table 1). While averaging over long proteins could lead to information loss in the resulting embeddings, we did not observe a correlation between performance and protein length (Fig. S2, Table S1). In order to obtain the embeddings, we processed query and lookup protein the same way. If those have similar function and similar length, their embeddings might have lost information in the same way. This loss might have “averaged out” to generate similar embeddings.

Including more neighbors (*k*>1) only slightly affected F_max_ (Table S2; all F_max_ averages for *k=2* to *k*=10 remained within the 95% confidence interval of that for *k*=1). When taking all predictions into account independent of a threshold in prediction strength referred to as the reliability index (RI, Methods; i.e. even low confidence annotations are transferred), the number of predicted GO terms increased with higher *k* (Table S3). The average number of GO terms annotated per protein in *GOA2017* already reached 37, 14, 9 for BPO, MFO, CCO, respectively. When including all predictions independent of their strength (RI) our method predicted more terms for CCO and BPO than expected from this distribution even for *k*=1. Only for MFO the average (11.7 terms) predicted was slightly lower than expected for *k*=1 (number of terms exploded for *k*>1: Table S3). While precision dropped with adding terms, recall increased (Table S3). To avoid overprediction and given that *k* hardly affected F_max_, we chose *k*=1. This choice might not be best in practice: considering more than one hit (*k*>1) might help when the closest hit only contains unspecific terms. However, such a distinction will be left to expert users.

When applying the <u>*LK* evaluation mode</u> (limited-knowledge, i.e. query already has some annotation about function, Methods/CAFA3), the embedding-based annotation transfer reached F_max_ scores of 49±1%, 52±2%, and 58±3% for BPO, MFO, and CCO, respectively (Fig. S3). Thus, the embedding-based annotation transfer reached higher values for proteins with prior annotations (LK evaluation mode) than for novel proteins without any annotations (NK evaluation mode; Table 1); the same was true for the CAFA3 top-10 for which the F_max_ scores increased even more than for our method for BPO and MFO, and less for CCO (Fig. 1, Fig. S3). In the *LK* mode, predictions are evaluated for proteins for which 1-2 GO ontologies had annotations while those for another ontology (or two) were added after the CAFA3 deadline^9,19^. While supervised training uses such labels; our approach did not since we had excluded all CAFA3 targets explicitly from the annotation transfer database (*GOA2017*). Thus, our method could not benefit from previous annotations, i.e. *LK* and *NK* should be identical. The observed differences were most likely explained by how F_max_ is computed. The higher F_max_ score, especially for BPO, might be explained by data set differences, e.g. LK average number of BPO annotations was 19 compared to 26 for NK. Other methods might have reached even higher by training on known annotations.

### Embedding-based transfer successfully identified distant relatives

Embeddings condense information learned from sequences; identical sequences produce identical embeddings: if PIDE(Q,T)=100%, then DIST^emb^(Q,T)=0 (Eqn. 4). We had assumed a simple relation: the more similar two sequences, the more similar their embeddings because the underlying LMs only use sequences as input. Nevertheless, we observed embedding-based annotation transfer to outperform (higher F_max_) sequence-based transfer (Table 1, Fig. 1). This suggested embeddings to capture information beyond raw sequences. Explicitly calculating the correlation between sequence and embedding similarity for 431,224 sequence pairs from CAFA3/*GOA2017*-100, we observed a correlation of ρ=0.29 (Spearman’s correlation coefficient, p-value < 2.2e-16; Table 2). Thus, sequence and embedding similarity correlated at an unexpectedly low level. However, our results demonstrated that embedding similarity identified more distant relatives than sequence similarity (Figs. S1, S4).

**Table 2:** Embedding and sequence similarity correlated^*^

| | *PIDE* | $d_{\text{SeqVec}}$ | $d_{\text{ProtBert}}$ |
| --- | --- | --- | --- |
| *PIDE* | 1 | 0.293 | 0.248 |
| $d_{\text{SeqVec}}$ | | 1 | 0.576 |
| $d_{\text{ProtBert}}$ | | | 1 |

\* **PIDE**: percentage pairwise sequence identity; $d_{\text{SeqVec}}$ : similarity in SeqVec<sup>20</sup> embeddings (Eqn. 5); $d_{\text{ProtBert}}$ : similarity in ProtBert<sup>21</sup> embeddings (Eqn. 5). All values represent Spearman's correlation coefficients calculated for 434,001 sequence pairs. For all pairs, the significance was $p\text{-value} < 2.2\text{e-}16$ , i.e. significant at the level of the precision of the software R<sup>28</sup>. The similarity between sequence and embeddings correlated less than the two different types of embeddings, namely SeqVec and ProtBert with each other. In order to highlight the trivial symmetry of the matrix, only the upper diagonal was given.

In order to quantify how different embeddings for proteins Q and T can still share GO terms, we redundancy reduced the *GOA2017* database used for annotation transfers at distance thresholds of decreasing PIDE with respect to the queries (in nine steps from 90% to 20%, Table S6). By construction, all proteins in *GOA2017-100* had PIDE<100% to all CAFA3 queries (Q). If the pair (Q,T) with the most similar embedding was also similar in sequence, embedding-based would equal sequence-based transfer. At lower PIDE thresholds, e.g. PIDE<20%, reliable annotation transfer through simple pairwise sequence alignment is no longer possible^14,29-32^. Although embeddings-based transfer tended to be slightly less successful for pairs with lower PIDE (Fig. 2: bars decrease toward right), the drop appeared small; on top, at almost all PIDE values, embedding-transfer remained above BLAST, i.e. sequence-based transfer (Fig. 2: most bars higher than reddish line - error bars show 95% confidence intervals). The exception was for MFO at PIDE<30% and PIDE<20% for which the F_max_ scores from sequence-based transfer (BLAST) were within the 95% confidence interval (Fig. 2). This clearly showed that our approach benefited from information available through embeddings but not through sequences, and that at least some protein pairs close in embedding and distant in sequence space might function alike. In order to correctly predict the next token, protein LMs have to learn complex correlations between residues as it is impossible to remember the multitude of all possible amino acid combinations in hundreds of millions to billions of protein sequences. This forces models to abstract higher level features from sequences. For instance, secondary structure can directly be extracted from embeddings through linear projection^26^. The LMs (SeqVec & ProtBert) might even have learned to find correlations between protein pairs diverged into the “midnight zone” sequence comparison in which sequence similarity becomes random^29,33^. Those cases are especially difficult to detect by the most successful search methods such as BLAST^34^ or MMseqs2^35^ relying on finding similar seeds missing at such diverged levels.

**Fig. 2:**
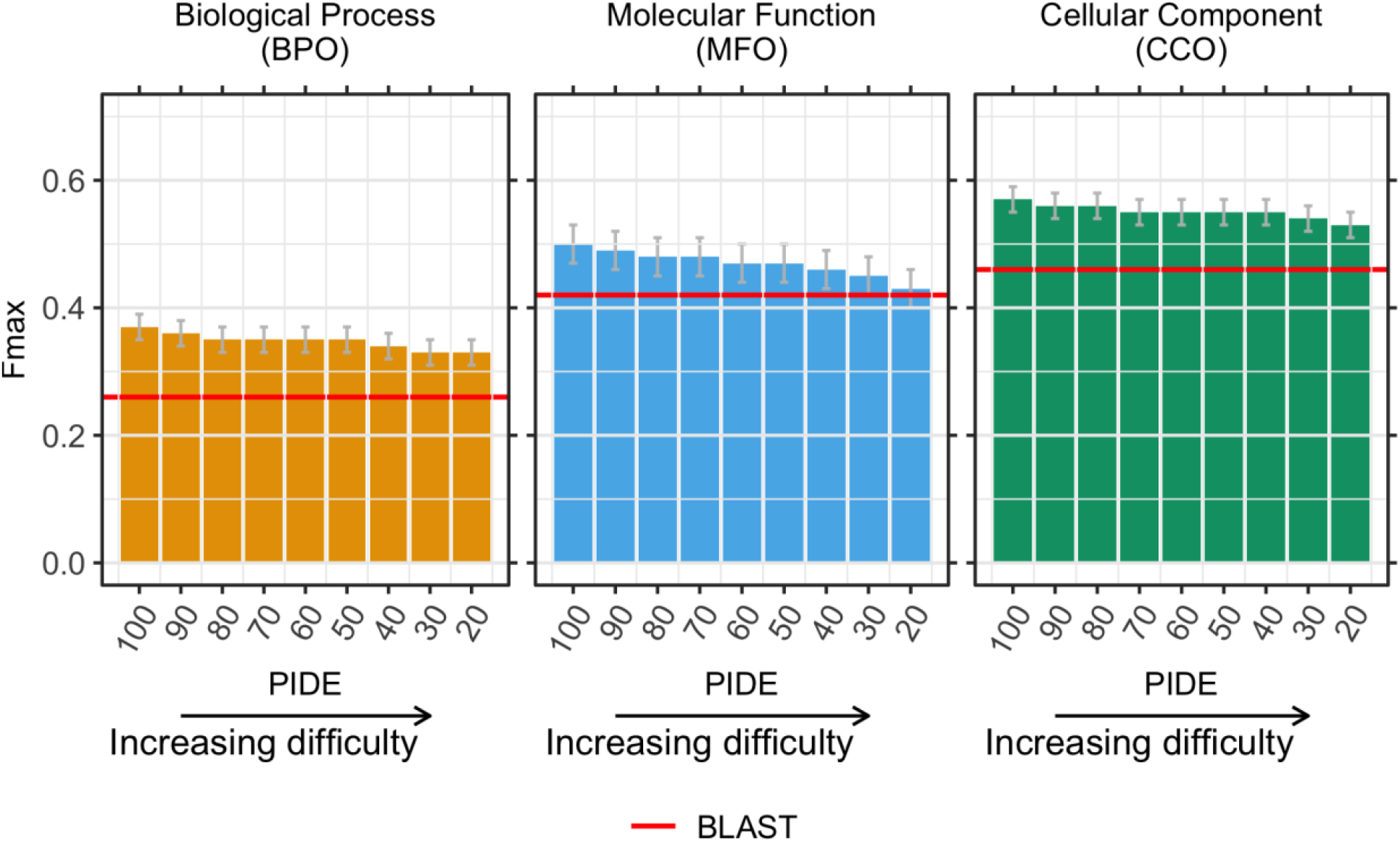
Embedding-based transfer succeeded for rather diverged proteins. After establishing the low correlation between embedding- and sequence-similarity, we tested how the level of percentage pairwise sequence identity (PIDE, x-axes) between the query (protein without known GO terms) and the transfer database (proteins with known GO terms, here subsets of *GOA2017)* affected the performance of the embedding-based transfer. Technically, we achieved this by removing proteins above a certain threshold in PIDE (decreasing toward right) from the transfer database. The y-axes showed the F_max_ score as compiled by CAFA3^19^. If embedding similarity and sequence identity correlated, our method would fall to the level of the reddish lines marked by BLAST. On the other hand, if the two were completely uncorrelated, the bars describing embedding-transfer would all have the same height (at least within the standard errors marked by the gray vertical lines at the upper end of each bar), i.e. embedding-based transfer would be completely independent of the sequence similarity between query and template (protein of known function). The observation that all bars tended to fall toward the right implied that embedding and sequence similarity correlated (although for CCO, F_max_ remained within the 95% confidence interval of F_max_ for *GOA2017-100).* The observation that our method remained mostly above the baseline predictions demonstrates that embeddings capture important orthogonal information. Error bars indicate 95% confidence intervals.

Ultimately, we failed to really explain why the abstracted level of sequences captured in embeddings outperformed raw sequences. One attempt at addressing this question led to displaying cases for which one of the two worked better (Fig. S5). Looking in more detail at outliers (embeddings more similar than sequences), we observed that embedding-based inference tended to identify more reasonable hits in terms of lineage or structure. For instance, for the uncharacterized transporter YIL166C (UniProt identifier P40445) from *Saccharomyces cerevisiae* (baker’s yeast), the closest hit in SeqVec embedding space was the high-affinity nicotinic acid transporter (P53322) also from *Saccharomyces cerevisiae.* Both proteins belong to the allantoate permease family while the most sequence-similar hit (with PIDE=31%) was the gustatory and odorant receptor 22 (Q7PMG3) from the insect *Anopheles gambiae* belonging to the gustatory receptor family. Experimental 3D structures were not available for any of the three proteins. However, comparative modeling using Swiss-Model^36^ revealed that both the target and the hit based on SeqVec were mapped to the same template (root-mean-square deviation (RMSD)=0.3Å) (Fig. S6a) while the hit based on sequence similarity was linked to a different structure (with RMSD=16.8Å) (Fig.S6b). Similarly, for the GDSL esterase/lipase At3g48460 (Q9STM6) from *Arabidopsis thaliana,* the closest hit in ProtBert embedding space was the GDSL esterase/lipase 2 (Q9SYF0) also from *Arabidopsis thaliana* while the most sequence-similar hit was the UDP-glucose 4-epimerase (Q564Q1) from *Caenorhabditis elegans.* The target and the embedding-based hit are both hydrolases belonging to the same CATH superfamily while the sequence-based hit is an isomerase and not annotated to any CATH superfamily. Comparative modeling suggested similar structures for target and embedding hit (RMSD=2.9Å) (Fig. S6c) while the structure found for the sequence-based hit similarity was very different (RMSD=26å) (Fig. S6d). This suggested embeddings to capture structural features better than just raw sequences. Homology-based inference depends on many parameters that can especially affect the resulting sequence alignment for distantly related proteins. Possibly, embeddings are more robust in identifying those more distant evolutionary relatives.

### Embedding-based transfer benefited from non-experimental annotations

Unlike the data set made available by CAFA3, annotations in our *GOA2017* data set were not limited to experimentally verified annotations. Instead, they included annotations inferred by computational biology, homology-based inference, or by “author statement evidence”, i.e. through information from publications. Using *GOA2017X,* the subset of *GOA2017-100* containing only experimental terms, our method reached F_max_=31±2% (P=28±2%, R=34±2%), 51±3% (P=53±3%, R=49±3%), and 56±2% (P=55±3%, R=57±3%) for BPO, MFO, and CCO, respectively. Compared to using *GOA2017-100,* the performance dropped significantly for BPO (F_max_=37±2% for *GOA2017-100,* Table 1); it decreased slightly (within 95% confidence interval) for CCO (F_max_=57±2% for *GOA2017-100,* Table 1); and it increased slightly (within 95% confidence interval) for MFO (F_max_=50±3% for *GOA2017-100,* Table 1). Thus, less reliable annotations might still help, in particular for BPO. Annotations for BPO may rely more on information available from publications that is not as easily quantifiable experimentally as annotations for MFO or CCO.

Many of the non-experimental annotations constituted sequence-based annotation transfers. Thus, non-experimental annotations might have helped because they constituted an implicit merger of sequence and embedding transfer. Adding homologous proteins might “bridge” sequence and embedding space by populating embedding space using annotations transferred from sequence space. The weak correlation between both spaces supported this speculation because protein pairs with very similar sequences may differ in their embeddings and *vice versa.*

### Improving annotations from 2017 to 2020 increased performance significantly

For CAFA3 comparisons, we only used data available before the CAFA3 submission deadline. When running new queries, annotations will be transferred from the latest GOA. We used *GOA2020-100* (from 02/2020 removing the CAFA3 targets) to assess how the improvement of annotations from 2017 to 2020 influenced annotation transfer (Table 1). On *GOA2020-100,* SeqVec embedding-based transfer achieved F_max_=50±2% (P=50±3%, R=50±3%), 60±3% (P=52±3%, R=71±3%), and 65±2% (P=57±3%, R=75±3%) for BPO, MFO, and CCO, respectively, for the *NK* evaluation mode (Table 1). This constituted a substantial increase over *GOA2017-100* (Table 1).

The large performance boost between *GOA2017* and *GOA2020* suggested the addition of many relevant GO annotations. However, for increasingly diverged pairs (Q,T), we observed a much larger drop in F_max_ than for *GOA2017* (Fig. 2, Fig. S4). In the extreme, *GOA2020-20* (PIDE(Q,T)<20%) with F_max_=33±2% (BPO), 44±2% (MFO), and 54±2% (CCO) fell to the same level as *GOA2017-20* (Fig. 2, Fig. S4). These results suggested that many of the relevant GO annotations were added for proteins sequence-similar to those with existing annotations. Put differently, many helpful new experiments simply refined previous computational predictions.

Running BLAST against *GOA2020-100* for sequence-based transfer (choosing the hit with the highest PIDE) showed that sequence-transfer also profited from improved annotations (difference in F_max_ values for BLAST in Table 1). However, while F_max_ scores for embedding-based transfer increased the most for BPO, those for sequence-based transfer increased most for MFO. Embedding-transfer still outperformed BLAST for the *GOA2020-100* set (Fig. S4c).

Even when constraining annotation transfer to sequence-distant pairs, our method outperformed BLAST against *GOA2020-100* in terms of F_max_ at least for BPO and for higher levels of PIDE in MFO/CCO (Fig. S4c). However, comparing the results for BLAST on the *GOA2020-100* set with the performance of our method for subsets of very diverged sequences (e.g. PIDE<40% for *GOA2020-40)* under-estimated the differences between sequence- and embedding-based transfer, because the two approaches transferred annotations from different data sets. For a more realistic comparison, we re-ran BLAST only considering hits below certain PIDE thresholds (for comparability we could not do this for CAFA3). As expected, performance for BLAST decreased with PIDE (Fig. S4 lighter bars), e.g. for PIDE<20%, F_max_ fell to 8% for BPO, 10% for MFO, and 11% for CCO (Fig. S4c lighter bars) largely due to low coverage, i.e. most queries had no hit to transfer annotations from. At this level (and for the same set), the embedding-based transfer proposed here, still achieved values of 33±2% (BPO), 44±2% (MFO), and 54±2% (CCO). Thus, our method made reasonable predictions at levels of sequence identity for which homology-based inference (BLAST) failed completely.

### Performance confirmed by new proteins

Our method and especially the threshold to transfer a GO term were “optimized” using the CAFA3 targets. Without any changes in the method, we tested a new data set of 298 proteins, *GOA2020-new,* with proteins for which experimental GO annotations have been added since the CAFA4 submission deadline (02/2020; Method). Using the thresholds optimized for CAFA3 targets (0.35 for BPO, 0.28 for MFO, 0.29 for CCO, Fig. 3), our method reached F1=50±11%, 54±5%, and 66±8% for BPO, MFO, and CCO, respectively. For BPO and CCO, the performance was similar to that for the CAFA3 targets; for MFO it was slightly below but within the 95% CI (Table 1). For yet a different set, submitted for MFO to CAFA4, the first preliminary evaluation published during ISMB2020^37^, also suggested our approach to make it to the top-ten, in line with the *post facto* CAFA3 results presented here.

**Fig. 3:**
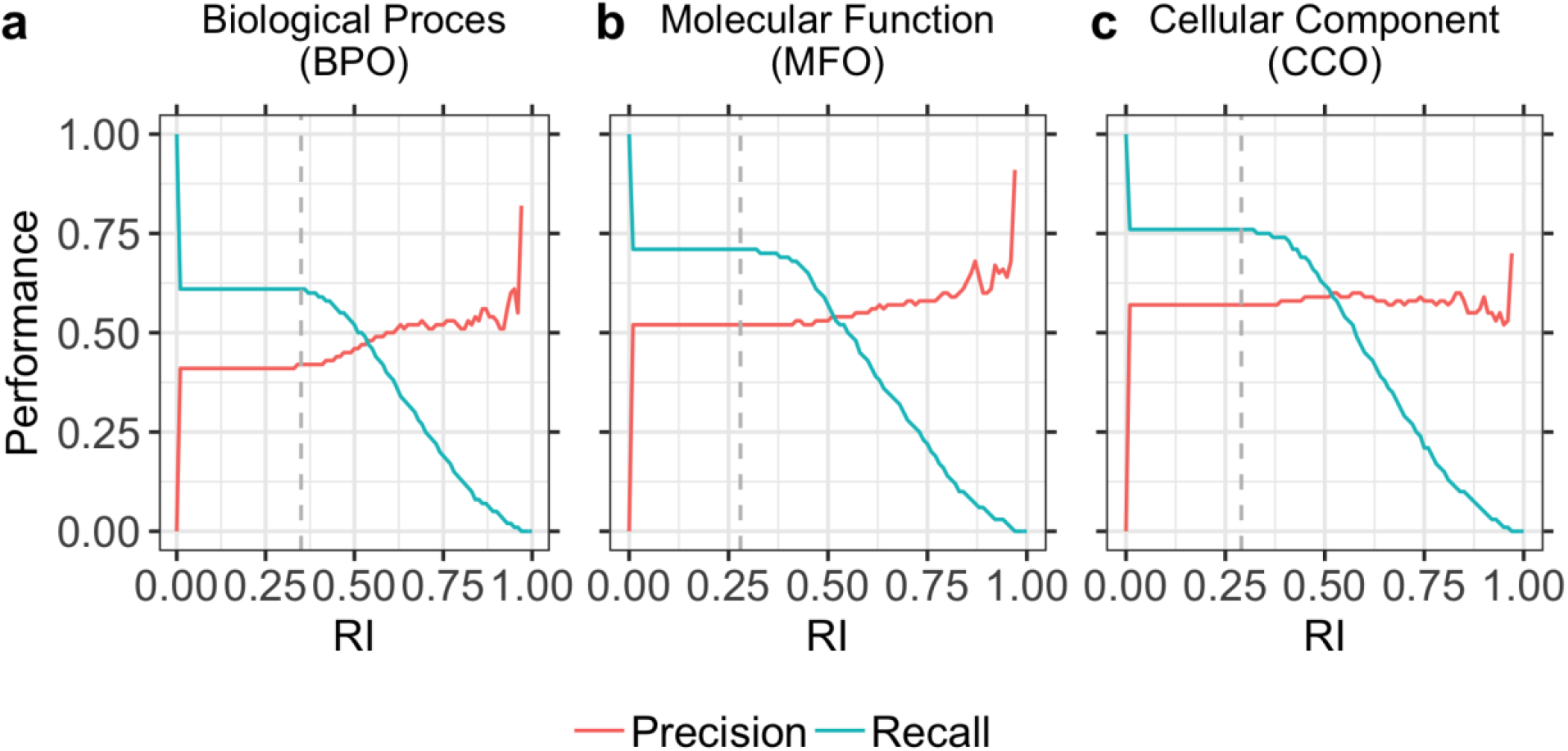
Precision and recall for different reliability indices (RIs). We defined a reliability index (RI) measuring the strength of a prediction (Eqn. 5), i.e. for the embedding proximity. Precision (Eqn. 1) and recall (Eqn. 2) were almost constant for lower RIs (up to ~ 0.3) for all three ontologies (BPO, MFO, CCO). For higher RIs, precision increased while recall dropped. However, due to the low number of terms predicted at very high RIs (>0.8), precision fluctuated and was not fully correlated with RI. Panel **a** shows precision and recall for BPO, panel **b** for MFO, and panel **c** for CCO. Dashed vertical lines marked the thresholds used by the CAFA3 tool to compute F_max_: 0.35 for BPO, 0.28 for MFO, and 0.29 for CCO. At least for BPO and MFO higher RI values correlated with higher precision, i.e. users could use the RI to estimate how good the predictions are likely to be for their query, or to simply scan only those predictions more likely to be correct (e.g. RI>0.8).

### Embedding similarity influenced performance

Homology-based inference works best for pairs with high PIDE. Analogously, we assumed embedding-transfer to be best for pairs with high embedding similarity, i.e. low Euclidean distance (Eqn. 4). We used this to define a reliability index (RI, Eqn. 5). For the *GOA2020-100* set, the minimal RI was 0.24. The CAFA evaluation determined 0.35 for BPO, 0.28 for MFO, and 0.29 for CCO as thresholds leading to optimal performance as measured by F_max_ (Fig. 3 dashed grey lines marked these thresholds). For all ontologies, precision and recall were almost constant for lower RIs (up to ~ 0.3). For higher RIs, precision increased, and recall decreased as expected (Fig. 3). While precision increased up to 82% for BPO, 91% for MFO, and 70% for CCO, it also fluctuated for high RIs (Fig. 3). This trend was probably caused by the low number of terms predicted at these RIs. For CCO, the RI essentially did not correlate with precision. This might point to a problem in assessing annotations for which the trivial Naïve method reached values of F_max_~55% outperforming most methods. Possibly, some prediction of the type “organelle” is all that is needed to achieve a high F_max_ in this ontology.

### Similar performance for different embeddings

We compared embeddings derived from two different language models (LMs). So far, we used embeddings from SeqVec^20^. Recently, ProtBert, a transformer-based approach using a masked language model objective (Bert^38^) instead of auto-regression and more protein sequences (BFD^39,40^) during pre-training, was shown to improve secondary structure prediction^21^. Replacing SeqVec by ProtBert embeddings to transfer annotations, our approach achieved similar F_max_ scores (Table 1). In fact, the ProtBert F_max_ scores remained within the 95% confidence intervals of those for SeqVec (Table 1). Similar results were observed when using *GOA2017-100* (Table 1).

On the one hand, the similar performance for both embeddings might indicate that both LMs extracted equally beneficial aspects of function, irrespective of the underlying architecture (LSTMs in SeqVec, transformer encoders in ProtBert) or training set (SeqVec used UniRef50 with ~33M proteins, ProtBert used BFD with ~2.1B proteins). On the other hand, the similar F_max_ scores might also highlight that important information was lost when averaging over the entire protein to render fixed-size vectors. The similarity in F_max_ scores was less surprising given the high correlation between SeqVec and ProtBert embeddings (ρ=0.58, p-value < 2.2e-16; Table 2). The two LMs correlated more with each other than either with PIDE (Table 2).

### No gain from simple combination of embedding- and sequence-based transfer

All three approaches toward annotation transfer (embeddings from SeqVec or ProtBert, and sequence) had strengths; although performing worse on average, for some proteins sequence-transfer performed better. In fact, analyzing the pairs for which embedding-based transfer or sequencebased transfer outperformed the other method by at least four percentage points (|F_max_(BLAST)- F_max_(embeddings)|≥4) illustrated the expected cases for which PIDE was high and embedding similarity low, and *vice versa,* along with more surprising cases for which low PIDE still yielded better predictions than relatively high embedding RIs (Fig. S5). Overall, these results (Fig. S5) again underlined that LM embeddings abstract information from sequence that are relevant for comparisons and not captured by sequences alone. However, it also indicates that even protein pairs with low embedding similarity can share similar GO terms. In fact, embedding similarity for SeqVec embeddings only weakly correlated with GO term similarity (Spearman rank coefficient (ρ=0.28, p-value < 2.2e-16), but proteins with identical GO annotations were on average more likely to be close than proteins with more different GO annotations (Fig. S7). The similarity of GO terms for two proteins was proxied through the Jaccard index (Eqn. 7). More details are provided in the SOM.

To benefit from the cases where BLAST outperformed our approach, we tried simple combinations: firstly, we considered all terms predicted by embeddings from either SeqVec or ProtBert. Secondly, reliability scores were combined leading to higher reliability for terms predicted in both approaches than for terms only predicted by one. None of those two improved performance (Table S4, method SeqVec/ProtBert). Other simple combinations also failed so far (Table S4, method SeqVec/ProtBert/BLAST). Future work might improve performance through more advanced combinations.

### Case study: embedding-based annotation transfer for three proteomes

Due to its simplicity and speed, embedding-based annotation transfer can easily be applied to novel proteins to shed light on their potential functionality. We applied our method to the proteomes of three different proteomes: human (20,370 proteins from Swiss-Prot) as a well-researched proteome, the fungus *Armillaria ostoyae* (22,192 proteins, 0.01% of these in Swiss-Prot), as one of the oldest (2500 years) and largest (4*10^5^ kg/spanning over 10 km^2^) living organisms known today^41^, and SARS-CoV-2, the virus causing COVID-19 (14 proteins). At RI=1.0, annotations were inferred from proteins of this organism (“self-hits”). Using only experimentally verified annotations (lookup data set *GOA2020X),* revealed both how few proteins were directly annotated (self-hits) in these organisms and how much of the sequence-annotation gap is gapped through embedding-based inference (Fig. 4: bars with darker orange, blue, green for BPO, CCO, and MFO respectively). In particular, for self-hits, i.e. proteins with 100% pairwise sequence identity (PIDE) to the protein with known annotation, it became obvious how few proteins in human have explicit experimental annotation (sum over all around 270), while through embedding-based inference up to 80% of all human proteins could be annotated through proteins from other organisms (light bars in Fig. 4 give results for the entire *GOA2020* which is dominated by annotations not directly verified by experiment). For the other two proteomes from the fungus *(Armillaria ostoyae)* and the coronavirus (SARS-CoV-2), there were no inferences at this high level. On the other end of including all inferences as assessed through the data presented in all other figures and tables (i.e. at the default thresholds), for all three proteins most proteins could be annotated directly from experimentally verified annotations through embeddings (three left-most bars in Fig. 4 for BPO, CCO, and MFO). In fact, when including all GO annotations from GOA (lookup set *GOA2020*), almost all proteins in all three proteomes could be annotated (Fig. 4: lighter colored left-most bars close to fraction of 1, i.e. all proteins). For SARS-CoV-2, our method reached 100% coverage (prediction for all proteins) already at RI≥0.5 (Fig. 4c, lighter colors, middle bars) through well-studied, similar viruses such as the human SARS coronavirus (SARS-CoV). RI=0.5 represent roughly a precision and recall of 50% for all three ontologies (Fig. 3). For *Armillaria ostoyae,* almost no protein was annotated through self-hits even when using unverified annotations (Fig. 4b: no bar at RI=1). At RI=0.5, about 25% of the proteins were annotated.

**Fig. 4:**
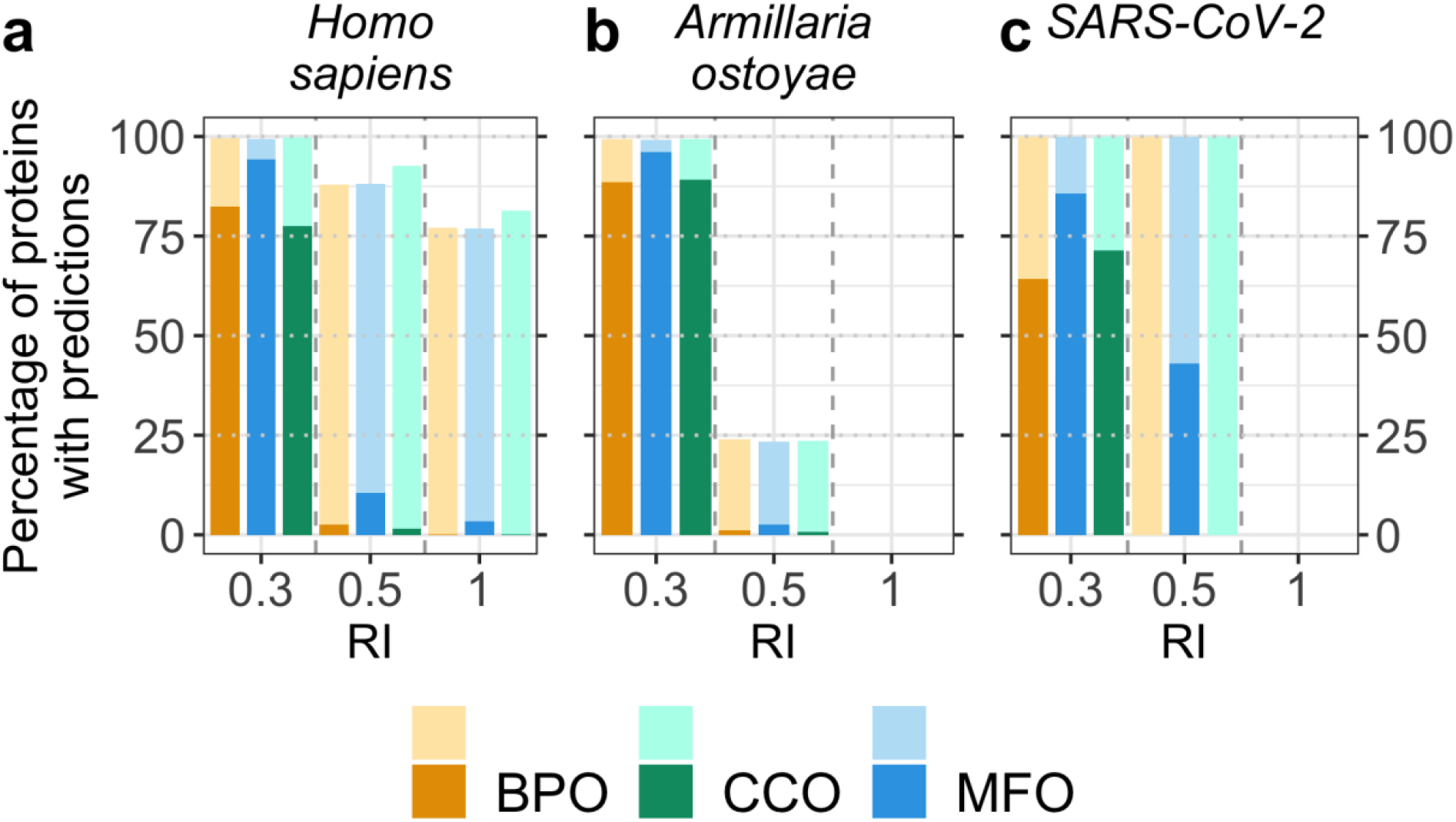
Fraction of proteomes with predicted GO terms. We applied our method to three proteomes (animal: *Homo sapiens,* fungus: *Armillaria ostoyae,* and virus: SARS-Cov-2) and monitored the fraction of proteins in each proteome for which our method predicted GO terms for different thresholds in embedding similarity (RI, Eqn. 5). We show predictions for RI=1.0 (“self-hits”), RI=0.5 (with an expected precision/recall=0.5), and RI=0.3 (CAFA3 thresholds). Darker colored bars indicate predictions using *GOA2020X* as lookup set (only experimentally verified GO annotations) and lighter colors indicate predictions using *GOA2020* as lookup set (using all annotations in GOA). **a.** The human proteome is well-studied (all 20,370 proteins are in Swiss-Prot) and for most proteins, GO annotations are available, but those annotations are largely not experimentally verified (very small, dark-colored bars vs large, lighter-colored bars at RI=1.0). **b.** The proteome of the fungus *Armillaria ostoyae* appears more exceptional (0.01% of the 22,192 proteins were in Swiss-Prot); at RI>=0.5, predictions could be made only for 25% of the proteins when also using unverified annotations and none of the proteins already had any GO annotations. **c.** While annotations were unknown for most proteins of the novel virus SARS-CoV-2 (no coverage at RI=1), many annotations could be transferred from the human SARS coronavirus (SARS-CoV) and the bat coronavirus HKU3 (BtCoV) allowing GO term predictions for all proteins at reliability values ≥0.5.

### Case study: embedding-based annotation transfer for SARS-CoV-2 proteome

Given the relevance of SARS-CoV-2, we did not only apply our method to predict GO terms (BPO, MFO, and CCO) for all 14 SARS-CoV-2 proteins (taken from UniProt^3^; all raw predictions were made available as additional files named predictions_$emb_$ont.txt replacing the variables $emb and $ont as follows: $emb=seqvec|protbert, and $ont=bpo|mfo|cco), but also investigated the resulting annotations further. While the two replicase polyproteins pp1a and pp1ab can also be split further into up to 12 non-structural proteins resulting in 28 proteins^42^, we used the definition from UniProt identifying 14 different proteins.

*Step 1: confirmation of known annotations.* Out of the 42 predictions (14 proteins in 3 ontologies), 12 were based on annotation transfers using proteins from the human SARS coronavirus (SARS-CoV), and 13 on proteins from the bat coronavirus HKU3 (BtCoV). CCO predictions appeared reasonable with predicted locations mainly associated with the virus (e.g. viral envelope, virion membrane) or the host (e.g. host cell Golgi apparatus, host cell membrane). Similarly, MFO predictions often matched well-known annotations, e.g. the replicase polyproteins 1a and 1ab were predicted to be involved in RNA-binding as confirmed by UniProt. In fact, annotations in BPO were known for 7 proteins (in total 40 GO terms), in MFO for 6 proteins (30 GO terms), and in CCO for 12 proteins (68 GO terms). Only three of these annotations were experimentally verified. With our method, we predicted 25 out of the 40 GO terms for BPO (63%), 14/30 for MFO (47%), and 59/68 for CCO (87%). Even more annotations were similar to the known GO annotations but were more or less specific (Table S5 summarized all predicted and annotated GO leaf terms, the corresponding names can be found in the additional files predictions_$emb_$ont.txt).

*Step 2: new predictions.* Since the GO term predictions matched well-characterized proteins, predictions might provide insights into the function of proteins without or with fewer annotations. For example, function and structure of the non-structural protein 7b (Uniprot identifier P0DTD8) are not known except for a transmembrane region of which the existence was supported by the predicted CCO annotation *“integral component of the membrane’’* and *“host cell membrane’.* This CCO annotation was also correctly predicted by the embedding-based transfer from an *Ashbya gossypii* protein. Additionally, we predicted *“transport of virus in host, cell to cell”* for BPO and *“proton transmembrane transporter activity’* for MFO. This suggested non-structural protein 7b to play a role in transporting the virion through the membrane into the host cell. Visualizing the leaf term predictions in the GO hierarchy could help to better understand very specific annotations. For the BPO annotation of the non-structural protein 7b, the tree revealed that this functionality constituted two major aspects: The interaction with the host and the actual transport to the host (Fig. S10). To visualize the predicted terms in the GO hierarchy, for example the tool NaviGO^43^ can be used which can help to interpret the GO predictions given for the SARS-CoV-2 proteins here.

Comparing annotation transfers based on embeddings from SeqVec and from ProtBert showed that 16 of the 42 predictions agreed for the two different language models (LMs). For five predictions, one of the two LMs yielded more specific annotations, e.g., for the nucleoprotein (Uniprot identifier P0DTC9) which is involved in viral genome packaging and regulation of viral transcription and replication. For this protein, SeqVec embeddings found no meaningful result, while ProtBert embeddings predicted terms such as *“RNA polymerase II preinitiation complex”* and *“positive regulation of transcription by RNA polymerase II”* fitting to the known function of the nucleoprotein. This example demonstrated how the combination of results from predictions using different LMs may refine GO term predictions.

## Conclusions

We introduce a new concept for the prediction of GO terms, namely the annotation transfer based on similarity of embeddings obtained from deep learning language models (LMs). This approach conceptually replaces sequence information by complex embeddings that capture some non-local information beyond sequence similarity. The underlying LMs (SeqVec & ProtBert) are highly involved and complex, and their training is time-consuming and data intensive. Once that is done, those pre-trained LMs can be applied, their abstracted understanding of the language of life as captured by protein sequences can be transferred to yield an extremely simple, yet effective novel method for annotation transfer. This novel prediction method complements homology-based inference. Despite its simplicity, this new method outperformed by several margins of statistically significance homology-based inference (“BLAST”) with F_max_ values of BPO **+11** ±2% (F_max_(embedding)-F_max_(sequence)), MFO **+8**±3%, and CCO **+11**±2% (Table 1, Fig. 1); it even might have reached the top ten, had it participated at CAFA3 (Fig. 1). Embedding-based transfer remained above the average for sequence-based transfer even for protein pairs with PIDE<20% (Fig. 2), i.e. embedding similarity worked for proteins that diverged beyond the recognition in pairwise alignments (Figs. S2 & S3). Embedding-based transfer is also blazingly fast to compute, i.e. around 0.05s per protein. The only time-consuming step is computing embeddings for all proteins in the lookup database which needs to be done only once; it took about 30min for the entire human proteome. GO annotations added from 2017 to 2020 improved both sequence- and embedding-based annotation transfer significantly (Table 1). Another aspect of the simplicity is that, at least in the context of the CAFA3 evaluation, the choice of none of the two free parameters really mattered: embeddings from both LMs tested performed, on average, equally, and the number of best hits (k-nearest neighbors) did not matter much (Table S2). The power of this new concept is generated by the degree to which embeddings implicitly capture important information relevant for protein structure and function prediction. One reason for the success of our new concept was the limited correlation between embeddings and sequence (Table 2). Additionally, the abstraction of sequence information in embeddings appeared to make crucially meaningful information readily available (Fig. S6). This implies that embeddings have the potential to revolutionize the way sequence comparisons are carried out.

## Methods

### Generating embedding space

The embedding-based annotation transfer introduced here requires each protein to be represented by a fixed-length vector, i.e. a vector with the same dimension for a protein of 30 and another of 40,000 residues (maximal sequence length for ProtBert). To this end, we used SeqVec^20^ to represent each protein in our data set by a fixed size embedding. SeqVec is based on ELMo^44^ using a stack of LSTMs^45^ for auto-regressive pre-training^46,47^ i.e. predicting the next token (originally a word in a sentence, here an amino acid in a protein sequence), given all previous tokens. Two independent stacks of LSTMs process the sequence from both directions. During pre-training, the two directions are joined by summing their losses; concatenating the hidden states of both directions during inference lets supervised tasks capture bi-directional context. For SeqVec, three layers, i.e. one uncontextualized CharCNN^48^ and two bi-directional LSTMs, were trained on each protein in UniRef50 (UniProt^3^ clustered at 50% PIDE resulting in ~33M proteins). In order to increase regularization, the weights of the token representation (CharCNN) as well as the final Softmax layer were shared between the two LSTM directions, and a 10% dropout rate was applied. For SeqVec, the CharCNN as well as each LSTM has a hidden state of size 512, resulting in a total of 93M free parameters. As only unlabeled data (no phenotypical data) was used (self-supervised training), the embeddings could not capture any explicit information such as GO numbers. Thus, SeqVec does not need to be retrained for subsequent prediction tasks using the embeddings as input. The hidden states of the pre-trained model are used to extract features. Corresponding to its hidden state size, SeqVec outputs for each layer and each direction a 512-dimensional vector; in this work, only the forward and backward passes of the first LSTM layer were extracted and concatenated into a matrix of size L * 1024 for a protein with L residues. While the auto-regressive pre-training only allowed to gather contextual information from either direction, the concatenation of the representations allowed our approach to benefit from bi-directional context. A fixed-size representation was then derived by averaging over the length dimension, resulting in a vector of size 1024 for each protein (Fig. S11). This simple way of information pooling (also called *global average pooling)* outperformed in many cases more sophisticated methods in NLP^49^ and showed competitive performance in bioinformatics for some tasks^20,21,26^. Based on experience from NLP^49,50^, we also investigated the effect of using a different pooling strategy, i.e. maximum pooling, to derive fixed size representations from SeqVec embeddings.

To evaluate the effect of using different LMs to generate the embeddings, we also used a transformer-based LM trained on protein sequences (ProtBert-BFD^21^, here simply referred to as *ProtBert).* ProtBert is based on the LM BERT^38^ (Bidirectional Encoder Representations from Transformers^51^) which processes sequential data through the self-attention mechanism^52^. Self-attention compares all tokens in a sequence to all others in parallel, thereby capturing long-range dependencies better than LSTMs. BERT also replaced ELMo’s auto-regressive objective by masked language modeling during pre-training, i.e. reconstructing corrupted tokens from the input, which enables to capture bi-directional context. ProtBert was trained with 30 attention layers, each having 16 attention heads with a hidden state size of 1024 resulting in a total of 420M free parameters which were optimized on 2.1B protein sequences (BFD)^39,40^ which is 70 times larger than UniRef50. The output of the last attention layer of ProtBert was used to derive a 1024-dimensional embedding for each residue. As for SeqVec, the resulting L * 1024 matrix was pooled by averaging over protein length providing a fixed-size vector of dimension 1024 for each protein. Usually, BERT’s special CLS-token is used for sequence-classification tasks^38^ as it is already optimized during pre-training on summarizing sequence information by predicting whether two sentences are consecutive in a document or not. In the absence of such a concept for proteins, this second loss was dropped from ProtBert’s pre-training rendering the CLS token without further fine-tuning on supervised tasks uninformative.

Embeddings derived from LMs change upon retraining the model with a different random seed, even using the same data and hyper-parameters. They are likely to change more substantially when switching the training data or tuning hyper-parameters. As retraining LMs is computationally (and environmentally) expensive, we leave assessing the impact of fine-tuning LMs to the future.

Generating the embeddings for all human proteins using both SeqVec and ProtBert allowed estimating the time required for the generation of the input to our new method. Using a single Nvidia GeForce GTX1080 with 8GB vRAM and dynamic batching (depending on the sequence length), this took, on average, about 0.05s per protein^21^.

### Data set

To create a database for annotation transfer, we extracted protein sequences with annotated GO terms from the Gene Ontology Annotation (GOA) database^53-55^ (containing 29,904,266 proteins from UniProtKB^3^ in February 2020). In order to focus on proteins known to exist, we only extracted records from Swiss-Prot^56^. Proteins annotated only at the ontology roots, i.e. proteins limited to “GO:0003674” (molecular_function), “GO:0008150” (biological_process), or “GO:0005575” (cellular_component) were considered meaningless and were excluded. The final data set *GOA2020* contained 295,558 proteins (with unique identifiers, IDs) described by 31,485 different GO terms. The GO term annotation for each protein includes all annotated terms and all their parent terms. Thereby, proteins are, on average, annotated by 37 terms in BPO, 14 in MFO, and 9 in CCO. Counting only leaves brought the averages to 3 in BPO, 2 in MFO, and 3 in CCO.

For comparison to methods that contributed to CAFA3^19^, we added another data set *GOA2017* using the GOA version available at the submission deadline of CAFA3 (Jan 17, 2017). After processing (as for *GOA2020*), *GOA2017* contained 307,287 proteins (unique IDs) described by 30,124 different GO terms. While we could not find a definite explanation for having fewer proteins in the newer database *(GOA2020* 295K proteins vs. *GOA2017* with 307K), we assume that it originated from major changes in GO including the removal of obsolete and inaccurate annotations and the refactoring of MFO^2^.

The above filters neither excluded GOA annotations inferred from phylogenetic evidence and author statements nor those based on computational analysis. We constructed an additional data set, *GOA2017X* exclusively containing proteins annotated in Swiss-Prot as experimental (evidence codes EXP, IDA, IPI, IMP, IGI, IEP, TAS, or IC) following the CAFA3 definition^19^. We further excluded all entries with PIDE=100% to any CAFA3 target bringing *GOA2017X* to 303,984 proteins with 28,677 different GO terms.

### Performance evaluation

The targets from the CAFA3 challenge^19^ were used to evaluate the performance of our new method. Of the 130,827 targets originally released for CAFA3, experimental GO annotations were obtained for 3,328 proteins at the point of the final benchmark collection in November 2017^19^. This set consisted of the following subsets with experimental annotations in each sub-hierarchy of GO: BPO 2,145, MFO 1,101, and CCO 1,097 (more details about the data set are given in the original CAFA3 publication^19^).

We used an additional data set, dubbed *GOA2020-new,* containing proteins added to GOA after February 2020, i.e. the point of accession for the GOA set used during the development of our method in preparation for CAFA4. This set consisted of 298 proteins with experimentally verified GO annotations and without any identical hits (i.e. 100% PIDE) in the lookup set *GOA2020*.

In order to expand the comparison of the transfer based on sequence- and embedding similarity, we also reduced the redundancy through applying CD-HIT and PSI-CD-HIT^57^ to the *GOA2020* and *GOA2017* sets against the evaluation set at thresholds *θ* of PIDE=100, 90, 80, 70, 60, 50, 40, 30 and 20% (Table S6 in the Supporting Online Material (SOM) for more details about these nine subsets).

We evaluated our method against two baseline methods used at CAFA3, namely *Naïve* and *BLAST,* as well as, against CAFA3’s top ten^19^. We computed standard performance measures. True positives (TP) were GO terms predicted above a certain reliability (RI) threshold (Method below), false positives (FP) were GO terms predicted but not annotated, and false negatives (FN) were GO terms annotated but not predicted. Based on these three numbers, we calculated precision (Eqn. 1), recall (Eqn. 2), and F1 score (Eqn. 3) as follows.

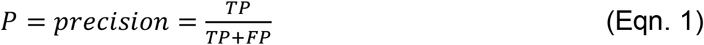

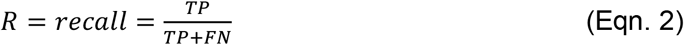

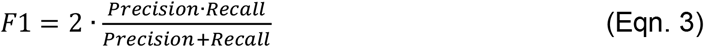

The F_max_ value denoted the maximum F1 score achievable for any threshold in reliability (RI, Eqn. 5). This implies that the assessment fixes the optimal value rather than the method providing this value. Although this arguably over-estimates performance, it has evolved to a quasi-standard of CAFA; the publicly available CAFA Assessment Tool^18,19^ calculated F_max_ for the CAFA3 targets in the same manner as for the official CAFA3 evaluation. If not stated otherwise, we reported precision and recall values for the threshold leading to F_max_.

CAFA3 assessed performance separately for two sets of proteins for all three ontologies: (i) proteins for which no experimental annotations were known beforehand (<u>no-knowledge, NK evaluation mode</u>) and (ii) proteins with some experimental annotations in one or two of the other ontologies (<u>limited-knowledge, LK evaluation mode</u>)^9,19^. We also considered these sets separately in our assessment. CAFA3 further distinguished between *full* and *partial evaluation* with *full evaluation* penalizing if no prediction was made for a certain protein, and *partial evaluation* restricting the assessment to the subset of proteins with predictions^19^. Our method predicted for every protein; thus, we considered only the *full evaluation.* Also following CAFA3, symmetric 95% confidence intervals were calculated as error estimates assuming a normal distribution and 10,000 bootstrap samples estimated mean and standard deviation.

### Method: annotation transfer through embedding similarity

For a given query protein Q, GO terms were transferred from proteins with known GO terms (sets *GOA2020* and *GOA2017)* through an approach similar to the k-nearest neighbor algorithm (k-NN)^58^. For the query Q and for all proteins in, e.g. *GOA2020,* the SeqVec^20^ embeddings were computed. Based on the Euclidean distance between two embeddings *n* and *m* (Eqn. 4), we extracted the *k* closest hits to the query from the database where *k* constituted a free parameter to optimize.

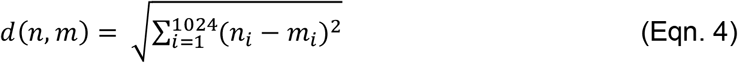

In contrast to standard k-NN algorithms, all annotations from all hits were transferred to the query instead of only the most frequent one^58^. When multiple pairs reached the same distance, all were considered, i.e. for a given *k*, more than *k* proteins might be considered for the GO term prediction. The calculation of the pairwise Euclidean distances between queries and all database proteins and the subsequent nearest neighbor extraction was accomplished very efficiently. For instance, the nearest-neighbor search of 1,000 query proteins against GOA20* with about 300,000 proteins took on average only about 0.005s per query on a single i7-6700 CPU, i.e. less than two minutes for all human proteins.

Converting the Euclidean distance enabled to introduce a reliability index (RI) ranging from 0 (weak prediction) to 1 (confident prediction) for each predicted GO term *p* as follows:

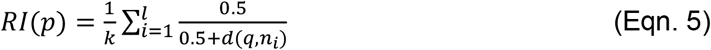

with *k* as the overall number of hits/neighbors, *l* as the number of hits annotated with the GO term *p* and the distance *d(q,n_i_)* between query and hit being calculated according to Eqn. 4.

Proteins represented by an embedding identical to the query protein (d=0) led to RI=1. Since the RI also takes into account, how many proteins *l* in a list of *k* hits are annotated with a certain term *p* (Eqn. 5), predicted terms annotated to more proteins (larger *l)* have a higher RI than terms annotated to fewer proteins (smaller *l*). As this approach accounts for the agreement of the annotations between the *k* hits, it requires the RI to be normalized by the number of considered neighbors *k*, making it not directly comparable for predictions based on different values for *k*. On top, if different embeddings are used to identify close proteins, RI values are not directly comparable, because embeddings might be on different scales.

Instead of assessing embedding proximity through the Euclidean distance, the embedding field typically uses the cosine distance (Eqn. 6):

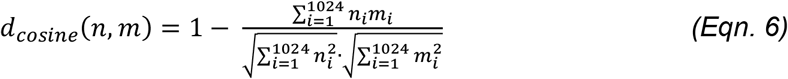

Our initial assessment suggested cosine and Euclidean distance to perform alike, and we chose to use the metric more familiar to structural biologists, namely the Euclidean distance throughout this analysis.

### GO term similarity

We measured the similarity between two sets of GO annotations *A* and *B* through the Jaccard index (Eqn. 7) where |*A ⋂ B*| is the number of GO terms present in both sets and |*A ⋃ B*| is the number of GO terms present in at least one of the sets (duplicates are only counted once):

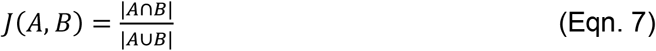

### Correlation analysis

We analyzed the correlation between sequence identity and embedding similarity through the Spearman’s rank correlation coefficient because our data was neither distributed normally, nor were the two measures for similarity measures linear. In contrast to, e.g. Pearson correlation, Spearman does not assume a normal distribution and detects monotonic instead of linear relations^59,60^.

### Availability

GO term predictions using embedding similarity for a certain protein sequence can be performed through our publicly available webserver: https://embed.protein.properties/. The source code along with all embeddings for *GOA2020* and *GOA2017*, and the CAFA3 targets are also available on GitHub: https://github.com/Rostlab/goPredSim (more details in the repository). In addition to reproducing the results, the source code also allows calculating embedding similarity using cosine distance.

## Supporting information

SARS-CoV-2 predictions

## Acknowledgements

Thanks to Tim Karl and Inga Weise (both TUM) for invaluable help with technical and administrative aspects of this work. Thanks to the organizers and participants of the CAFA challenge for their crucial contributions, in particular to Predrag Radivojac (Northeastern Boston), Iddo Friedberg (Iowa State Ames), Sean Mooney (U of Washington Seattle), Casey Green (U Penn Philadelphia, USA), Mark Wass (Kent, England), and Kimberly Reynolds (U Texas Dallas). Particular thanks to the maintainers of the Gene Ontology (in particular to Michael Ashburner - Cambridge and Suzanna Lewis - Berkeley), as well as, the Gene Ontology Annotation database (maintained by the team of Claire O’Donovan at the EBI) for their standardized vocabulary and curated data sets. Last, but not least, thanks to all maintainers of public databases and to all experimentalists who enabled this analysis by making their data publicly available. This work was supported by the Bavarian Ministry of Education through funding to the TUM, by a grant from the Alexander von Humboldt foundation through the German Ministry for Research and Education (BMBF: Bundesministerium für Bildung und Forschung), by a grant from BMBF (Software Campus, 01IS17049) as well as by a grant from Deutsche Forschungsgemeinschaft (DFG-GZ: RO1320/4-1). We gratefully acknowledge the support of NVIDIA Corporation with the donation of one Titan GPU used for this research.

## Data Availability

The source code and the embedding sets for target proteins and lookup databases are publicly available as a GitHub repository. GO term predictions for the SARS-CoV-2 proteins are provided as additional files.

## Abbreviations used

BERT: Bidirectional Encoder Representations from Transformers (particular deep learning language model)
BP(O): biological process (ontology) from GO
CAFA: Critical Assessment of protein Function Annotation algorithms
CC(O): cellular component (ontology) from GO
ELMo: Embeddings from Language Models
GO: gene ontology
GOA: Gene Ontology Annotation
k-NN: k-nearest neighbor
LK: limited-knowledge
LM: language model
LSTMs: Long-Short-Term-Memory cells
M: million
MF(O): molecular function (ontology) from GO
NK: no-knowledge
PIDE: percentage pairwise sequence identity
RI: reliability index
RMSD: Root-mean-square deviation

## Notes

https://github.com/Rostlab/goPredSim

https://embed.protein.properties/

### Competing Interest Statement

The authors have declared no competing interest.

## References

1 Krebs, H. A. & Johnson, W. A. Metabolism of ketonic acids in animal tissues. Biochem J 31, 645–660, doi:10.1042/bj0310645 (1937).

2 The Gene Ontology Consortium. The Gene Ontology Resource: 20 years and still GOing strong. Nucleic Acids Res 47, D330–D338, doi:10.1093/nar/gky1055 (2019).

3 UniProt Consortium. UniProt: a worldwide hub of protein knowledge. Nucleic Acids Res 47, D506–D515, doi:10.1093/nar/gky1049 (2019).

4 Hirst, J. D. & Sternberg, M. J. E. Prediction of structural and functional features of protein and nucleic acid sequences by artificial neural networks. Biochemistry 31, 615–623 (1992).

5 Rost, B., Liu, J., Nair, R., Wrzeszczynski, K. O. & Ofran, Y. Automatic prediction of protein function. Cellular and Molecular Life Sciences 60, 2637–2650 (2003).

6 Leslie, C., Eskin, E., Weston, J. & Noble, W. S. Mismatch string kernels for SVM protein classification. Bioinformatics, in press (2003).

7 Ofran, Y., Punta, M., Schneider, R. & Rost, B. Beyond annotation transfer by homology: novel protein-function prediction methods to assist drug discovery. Drug Discovery Today 10, 1475–1482 (2005).

8 Hamp, T. et al. Homology-based inference sets the bar high for protein function prediction. BMC Bioinformatics 14 Suppl 3, S7, doi:10.1186/1471-2105-14-S3-S7 (2013).

9 Radivojac, P. et al. A large-scale evaluation of computational protein function prediction. Nat Methods 10, 221–227, doi:10.1038/nmeth.2340 (2013).

10 Cozzetto, D., Minneci, F., Currant, H. & Jones, D. T. FFPred 3: feature-based function prediction for all Gene Ontology domains. Sci Rep 6, 31865, doi:10.1038/srep31865 (2016).

11 Kulmanov, M. & Hoehndorf, R. DeepGOPlus: improved protein function prediction from sequence. Bioinformatics 36, 422–429, doi:10.1093/bioinformatics/btz595 (2020).

12 Zuckerkandl, E. Evolutionary processes and evolutionary noise at the molecular level. J Mol Evol 7, 269–311 (1976).

13 Nakai, K. & Horton, P. PSORT: a program for detecting sorting signals in proteins and predicting their subcellular localization. Trends Biochem Sci 24, 34–36 (1999).

14 Nair, R. & Rost, B. Sequence conserved for sub-cellular localization. Protein Science 11, 2836–2847 (2002).

15 Goldberg, T. et al. LocTree3 prediction of localization. Nucleic Acids Res 42, W350–355, doi:10.1093/nar/gku396 (2014).

16 Qiu, J. et al. ProNA2020 predicts protein-DNA, protein-RNA, and protein-protein binding proteins and residues from sequence. J Mol Biol 432, 2428–2443, doi:10.1016/j.jmb.2020.02.026 (2020).

17 Goldberg, T., Rost, B. & Bromberg, Y. Computational prediction shines light on type III secretion origins. Sci Rep 6, 34516, doi:10.1038/srep34516 (2016).

18 Jiang, Y. et al. An expanded evaluation of protein function prediction methods shows an improvement in accuracy. Genome Biol 17, 184, doi:10.1186/s13059-016-1037-6 (2016).

19 Zhou, N. et al. The CAFA challenge reports improved protein function prediction and new functional annotations for hundreds of genes through experimental screens. Genome Biol 20, 244, doi:10.1186/s13059-019-1835-8 (2019).

20 Heinzinger, M. et al. Modeling aspects of the language of life through transfer-learning protein sequences. BMC Bioinformatics 20, 723, doi:10.1186/s12859-019-3220-8 (2019).

21 Elnaggar, A. et al. ProtTrans: towards cracking the language of life’s code through selfsupervised deep learning and high performance computing. bioRxiv (2020).

22 Mikolov, T., Cheng, K., Corrado, G. & Dean, J. Efficient Estimation of Word Representations in Vector Space in 1st International Conference on Learning Representations, ICLR 2013, Scottsdale, Arizona, USA, May 2-4, 2013, Workshop Track Proceedings.

23 Allen, C. & Hospedales, T. Analogies Explained: Towards Understanding Word Embeddings in Proceedings of the 36th International Conference on Machine Learning. 223–231 (PMLR).

24 Brokos, G.-I., Malakasiotis, P. & Androutsopoulos, I. Using Centroids of Word Embeddings and Word Mover’s Distance for Biomedical Document Retrieval in Question Answering in Proceedings of the 15th Workshop on Biomedical Natural Language Processing. 114–118 (Association for Computational Linguistics).

25 Kusner, M. J., Sun, Y., Kolkin, N. I. & Weinberger, K. Q. From Word Embeddings to Document Distances in Proceedings of the 32nd International Conference on International Conference on Machine Learning - Volume 37.

26 Rives, A. et al. Biological structure and function emerge from scaling unsupervised learning to 250 million protein sequences. bioRxiv, 622803, doi:10.1101/622803 (2020).

27 Vig, J. et al. BERTology Meets Biology: Interpreting Attention in Protein Language Models. arXiv (2020).

28 R Core Team. (R Foundation for Statistical Computing, 2017).

29 Rost, B. Twilight zone of protein sequence alignments. Protein Engineering 12, 85–94 (1999).

30 Rost, B. Enzyme function less conserved than anticipated. Journal of Molecular Biology 318, 595–608 (2002).

31 Mika, S. & Rost, B. Protein-protein interactions more conserved within species than across species. PLoS Computational Biology 2, e79, doi:10.1371/journal.pcbi.0020079 (2006).

32 Clark, W. T. & Radivojac, P. Analysis of protein function and its prediction from amino acid sequence. Proteins 79, 2086–2096, doi:10.1002/prot.23029 (2011).

33 Rost, B. Protein structures sustain evolutionary drift. Folding & Design 2, S19–S24 (1997).

34 Altschul, S. F., Gish, W., Miller, W., Myers, E. W. & Lipman, D. J. Basic local alignment search tool. J Mol Biol 215, 403–410, doi:10.1016/S0022-2836(05)80360-2 (1990).

35 Steinegger, M. & Söding, J. MMseqs2 enables sensitive protein sequence searching for the analysis of massive data sets. Nat Biotechnol 35, 1026–1028, doi:10.1038/nbt.3988 (2017).

36 Waterhouse, A. et al. SWISS-MODEL: homology modelling of protein structures and complexes. Nucleic Acids Res 46, W296–W303, doi:10.1093/nar/gky427 (2018).

37 El-Mabrouk, N. & Slonim, D. K. ISMB 2020 proceedings. Bioinformatics 36, i1–i2, doi:10.1093/bioinformatics/btaa537 (2020).

38 Devlin, J., Chang, M.-W., Lee, K. & Toutanova, K. BERT: Pre-training of Deep Bidirectional Transformers for Language Understanding in Proceedings of the 2019 Conference of the North American Chapter of the Association for Computational Linguistics: Human Language Technologies, Volume 1 (Long and Short Papers). 4171–4186 (Association for Computational Linguistics).

39 Steinegger, M., Mirdita, M. & Söding, J. Protein-level assembly increases protein sequence recovery from metagenomic samples manyfold. Nat Methods 16, 603–606, doi:10.1038/s41592-019-0437-4 (2019).

40 Steinegger, M. & Söding, J. Clustering huge protein sequence sets in linear time. Nat Commun 9, 2542, doi:10.1038/s41467-018-04964-5 (2018).

41 Anderson, J. B. et al. Clonal evolution and genome stability in a 2500-year-old fungal individual. Proc Biol Sci 285, 2018–2233, doi:10.1098/rspb.2018.2233 (2018).

42 O’Donoghue, S. I. et al. SARS-CoV-2 structural coverage map reveals state changes that disrupt host immunity. bioRxiv (2020).

43 Wei, Q., Khan, I. K., Ding, Z., Yerneni, S. & Kihara, D. NaviGO: interactive tool for visualization and functional similarity and coherence analysis with gene ontology. BMC Bioinformatics 18, 177, doi:10.1186/s12859-017-1600-5 (2017).

44 Peters, M. E. et al. Deep Contextualized Word Representations in Proceedings of the 2018 Conference of the North American Chapter of the Association for Computational Linguistics: Human Language Technologies, Volume 1 (Long Papers). 2227–2237 (Association for Computational Linguistics).

45 Hochreiter, S. & Schmidhuber, J. Long short-term memory. Neural Comput 9, 1735–1780, doi:10.1162/neco.1997.9.8.1735 (1997).

46 Mousa, A. & Schuller, B. Contextual Bidirectional Long Short-Term Memory Recurrent Neural Network Language Models: A Generative Approach to Sentiment Analysis in Proceedings of the 15th Conference of the {E}uropean Chapter of the Association for Computational Linguistics: Volume 1, Long Papers. 1023–1032 (Association for Computational Linguistics).

47 Peters, M., Ammar, W., Bhagavatula, C. & Power, R. Semi-supervised sequence tagging with bidirectional language models in Proceedings of the 55th Annual Meeting of the Association for Computational Linguistics (Volume 1: Long Papers). 1756–1765 (Association for Computational Linguistics).

48 Kim, Y., Jernite, Y., Sontag, D. & Rush, A. M. Character-Aware Neural Language Models in Proceedings of the Thirtieth AAAI Conference on Artificial Intelligence. (AAAI Press).

49 Shen, D. et al. Baseline Needs More Love: On Simple Word-Embedding-Based Models and Associated Pooling Mechanisms in Proceedings of the 56th Annual Meeting of the Association for Computational Linguistics (Volume 1: Long Papers). 440–450 (Association for Computational Linguistics).

50 Conneau, A., Douwe, K., Schwenk, H., Barrault, L. & Bordes, A. Supervised Learning of Universal Sentence Representations from Natural Language Inference Data in Proceedings of the 2017 Conference on Empirical Methods in Natural Language Processing. 670–680 (Association for Computational Linguistics).

51 Vaswani, A. et al. Attention is All you Need in Neural Information Processing Systems Conference. (eds I Guyon et al.) 5998–6008 (Curran Associates, Inc.).

52 Bahdanau, D., Cho, K. H. & Bengio, Y. Neural Machine Translation by Jointly Learning to Align and Translate in arXiv.

53 Camon, E. et al. The Gene Ontology Annotation (GOA) Database: sharing knowledge in Uniprot with Gene Ontology. Nucleic Acids Res 32, D262–266, doi:10.1093/nar/gkh021 (2004).

54 Huntley, R. P. et al. The GOA database: gene Ontology annotation updates for 2015. Nucleic Acids Res 43, D1057–1063, doi:10.1093/nar/gku1113 (2015).

55 GOA, http://www.ebi.ac.uk/GOA> (2020).

56 Boutet, E., Lieberherr, D., Tognolli, M., Schneider, M. & Bairoch, A. UniProtKB/Swiss-Prot. Methods Mol Biol 406, 89–112, doi:10.1007/978-1-59745-535-0_4 (2007).

57 Fu, L., Niu, B., Zhu, Z., Wu, S. & Li, W. CD-HIT: accelerated for clustering the nextgeneration sequencing data. Bioinformatics 28, 3150–3152, doi:10.1093/bioinformatics/bts565 (2012).

58 Cover, T. & Hart, P. Nearest neighbor pattern classification. IEEE Transactions on Information Theory 13, 21–27 (1967).

59 Dodge, Y. in The Concise Encyclopedia of Statistics 502–505 (Springer New York, 2008).

60 Spearman, C. The Proof and Measurement of Association Between Two Things. American Journal of Psychology 15, 72–101 (1904).

