## Supporting Online Material for "Embeddings from deep learning transfer GO annotations beyond homology"

### Supporting online material (SOM) for: Embeddings from deep learning transfer GO annotations beyond homology

### Table of Contents for Supporting Online Material (SOM)

### Short description of Supporting Online Material

In this Supporting Online Material (SOM), we show a more detailed performance assessment of our method to predict GO terms and provide details regarding the used data sets as well as a sketch of how to extract embeddings via SeqVec (Fig. S11). While averaging over long proteins could lead to information loss in the resulting embeddings, the performance of our method did not correlate with the protein length (Fig. S2, Table S1).

As $F_{max}$, precision and recall decreased for lookup sets redundancy reduced at lower sequence identity thresholds for *GOA2017* and *GOA2020* (Fig. S1, S4). Performance generally increased for the top ten CAFA3 competitors as well as for our method for the *LK* evaluation mode (Fig. S3). In the *LK* evaluation mode, targets are evaluated for which some annotations were already known at point of submission for CAFA3 and that gained additional annotations since then. Comparing the performance of homology-based inference with our method did not show a clear correlation between embedding similarity, sequence identity, and which of the methods performed better (Fig. S5). A more detailed analysis of two example targets and the respective hits through embedding- and sequence similarity shows that embeddings seem to better capture structural relationship (Fig. S6). We also evaluate our underlying assumption that proteins close in embedding space should have similar GO annotations (Fig. S7).

We applied our method to three different organisms (human, the fungus *Armillaria ostoyae*, and virus: SARS-CoV-2) and show the fraction of proteins with a GO term prediction for different RI thresholds using *GOA2020* (Fig. S8) or *GOA2020X* (Fig. S9). For further analysis of the predictions, the predicted leaf terms can be visualized in the GO tree structure. Fig. S10 shows an example for the non-structural protein 7b of SARS-CoV-2.

The choice of neighbors (*k*) to include for annotation transfer did not affect $F_{max}$ (Table S2), however including more hits than the closest one could increase recall and the quality of predictions if e.g. only unspecific terms are annotated to the closest hit (Table S3). A detailed analysis of annotated and predicted GO terms for the 14 proteins from SARS-CoV-2 showed that only three of the annotated GO terms are experimentally verified and that our method predicted 83%, 47%, and 87% of the annotated terms for BPO, MFO, and CCO (Table S4). Surprisingly, the *GOA2017* set was larger than *GOA2020* in terms of sequences (Table S1), however not in terms of GO annotations. In general, the redundancy-reduced versions of the two sets had on average a sequence similarity of 43-44% while the number of identical sequences (by identifier) dropped for the sets reduced at 20 and 30% sequence identity compared to the other sets (Table S5).

### Material

#### Fig. S1: Precision and recall for different lookup sets based on *GOA2017*

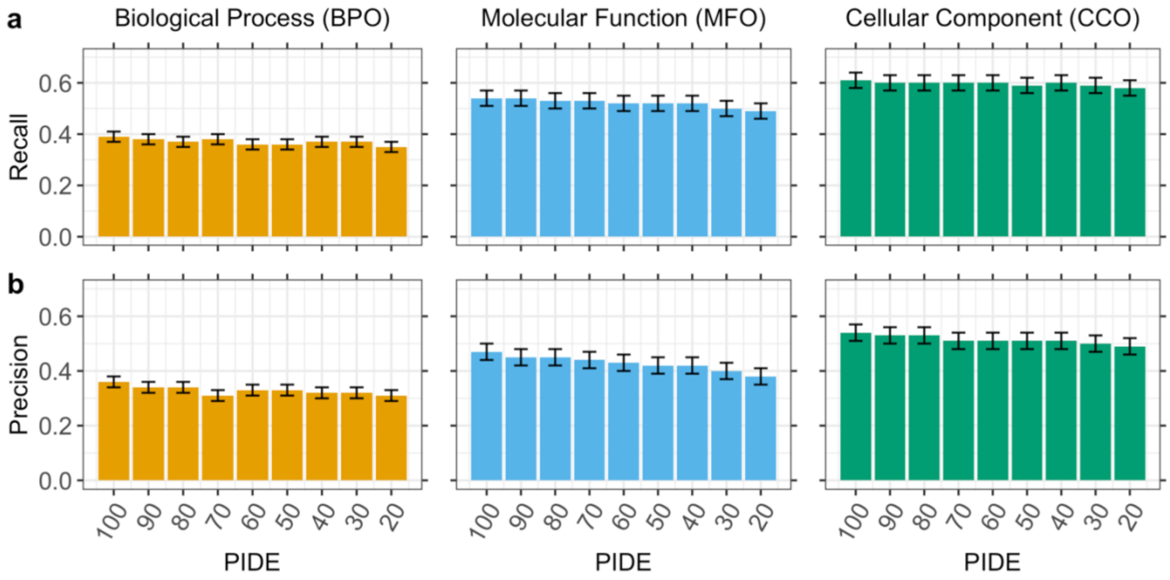

To test how the level of percentage pairwise sequence identity affects the performance of our method, we removed proteins above a certain pairwise sequence identity (as indicated on the x-axes) to the targets from our lookup set based on GOA version 2017. Panel **a** shows the recall and panel **b** the precision for BPO, MFO, and CCO, respectively, for lookup sets of <100, 90, 80, 70, 60, 50, 40, 30, and 20% sequence identity to the target proteins. Error bars indicate 95% confidence intervals.

#### Fig. S2: $F_{max}$ of our method for different protein lengths

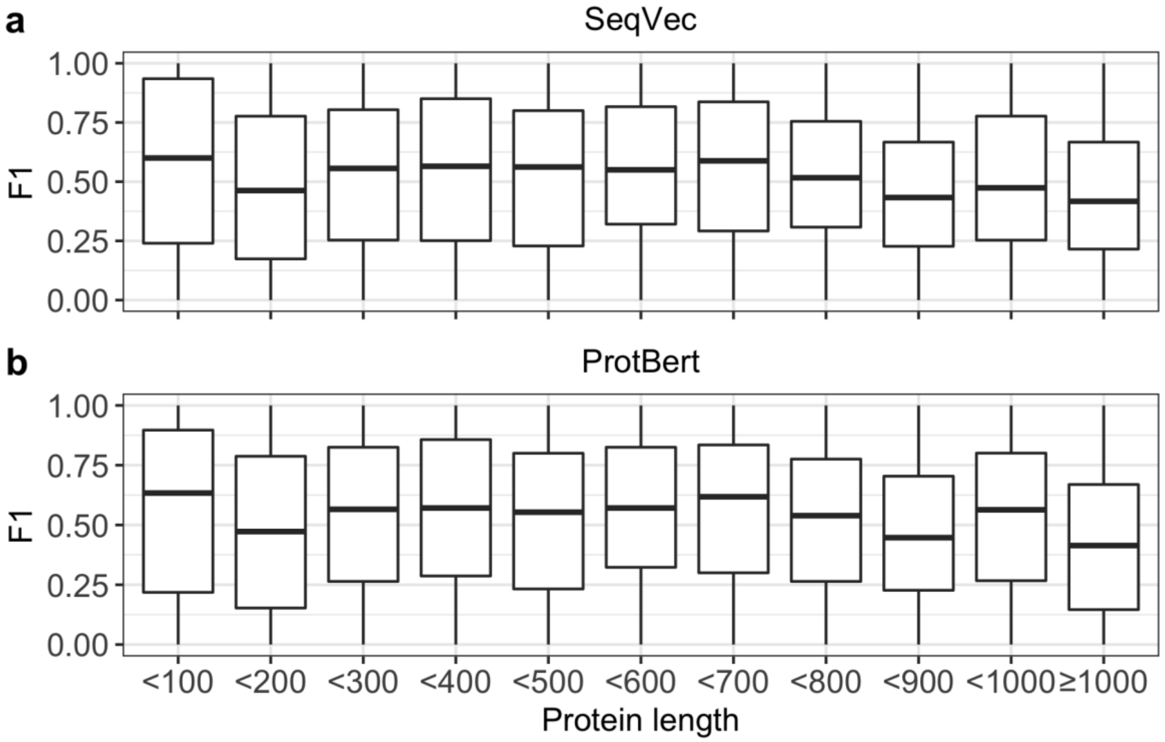

**Fig. S2: Performance not correlated with protein lengths.** We show the $F_{max}$ score for different intervals of varying protein length ranging from 100 to 1000 in steps of 100 for **a.** SeqVec and **b.** ProtBert. Protein embeddings for both models are derived by global average pooling over per-residue representations. This process could lead to an information loss for longer proteins. However, the performance of our method did not correlate with the length of the query protein.

#### Fig. S3: $F_{max}$ for our method and CAFA3 competitors using LK evaluation mode

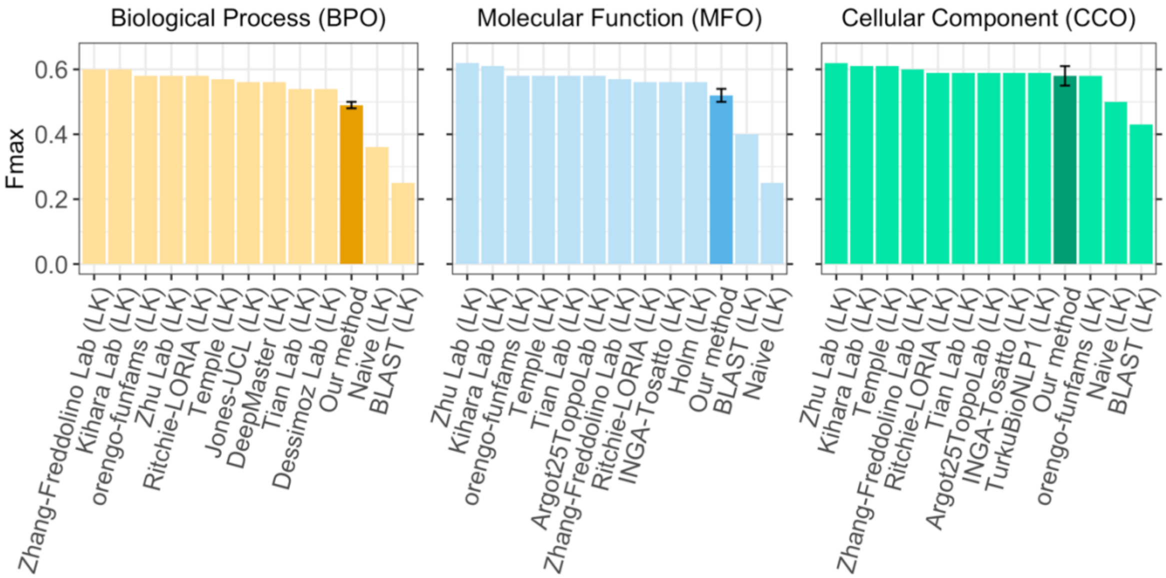

We compared the $F_{max}$ of our method (dark bar) for the three ontologies (BPO, MFO, CCO) to the top ten methods that did actually compete at CAFA3 [1] and to two background approaches (lighter bars) also applied in the CAFA3 challenge for the *LK* evaluation mode using proteins for which some GO terms have been known before.

#### Fig. S4: $F_{max}$, precision, and recall for different lookup sets based on *GOA2020*

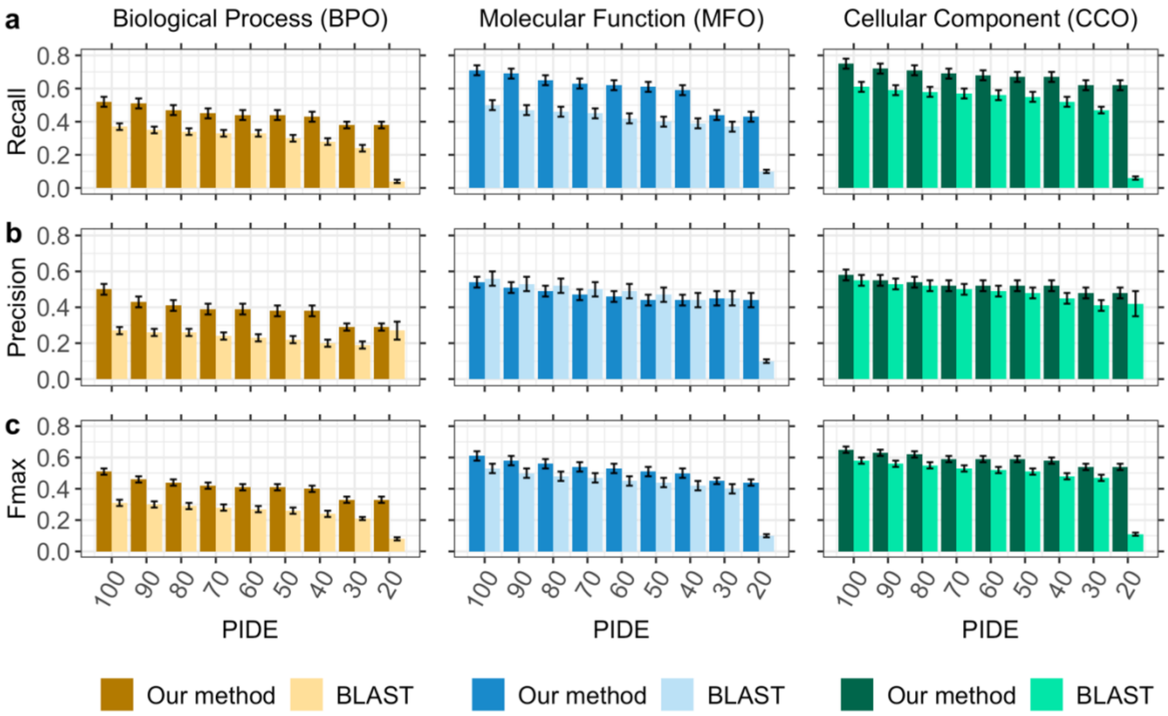

To test how the level of percentage pairwise sequence identity (PIDE) affects the performance of our method and of homology-based inference (BLAST), we stepwise removed proteins from our lookup set (here: GOA-2020) if they shared more than a certain PIDE (as indicated on the x-axes) to any query protein. Panel **a** shows the recall, panel **b** the precision, and panel **c** $F_{max}$ for BPO, MFO, and CCO, respectively, for lookup sets of <100, 90, 80, 70, 60, 50, 40, 30, and 20% PIDE to the target proteins. Sequence-based transfer was accomplished by running BLAST against *GOA2020* and transferring annotations from the hit with the highest PIDE (and PIDE <x% for the different lookup sets) in the local alignment (result marked by lighter colored bars, labeled as *BLAST*). Error bars indicate 95% confidence intervals.

**Fig. S5: Proteins for which embedding-based or homology-based inference worked better dependent of RI and PIDE**

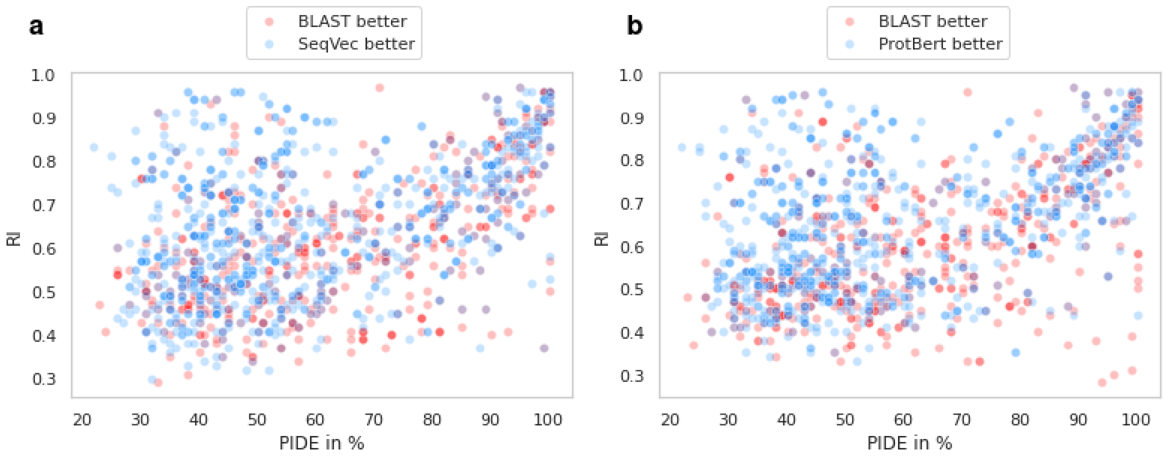

Blue points indicate proteins for which embedding-based annotation transfer (**a.** SeqVec, **b.** ProtBert) worked better than homology-based inference (BLAST) (by at least four percentage points), red points indicate the reverse: proteins for which PIDE worked better than embeddings. The RI gives the embedding similarity, PIDE is percentage pairwise sequence identity. There is no clear relationship between PIDE, RI, and which of the two approaches worked better. Only for high RIs and low PIDE, embedding-based annotation transfer performed almost always better than PIDE.

#### Fig. S6: Comparative model for two target proteins and the corresponding hits found through embedding- or sequence similarity

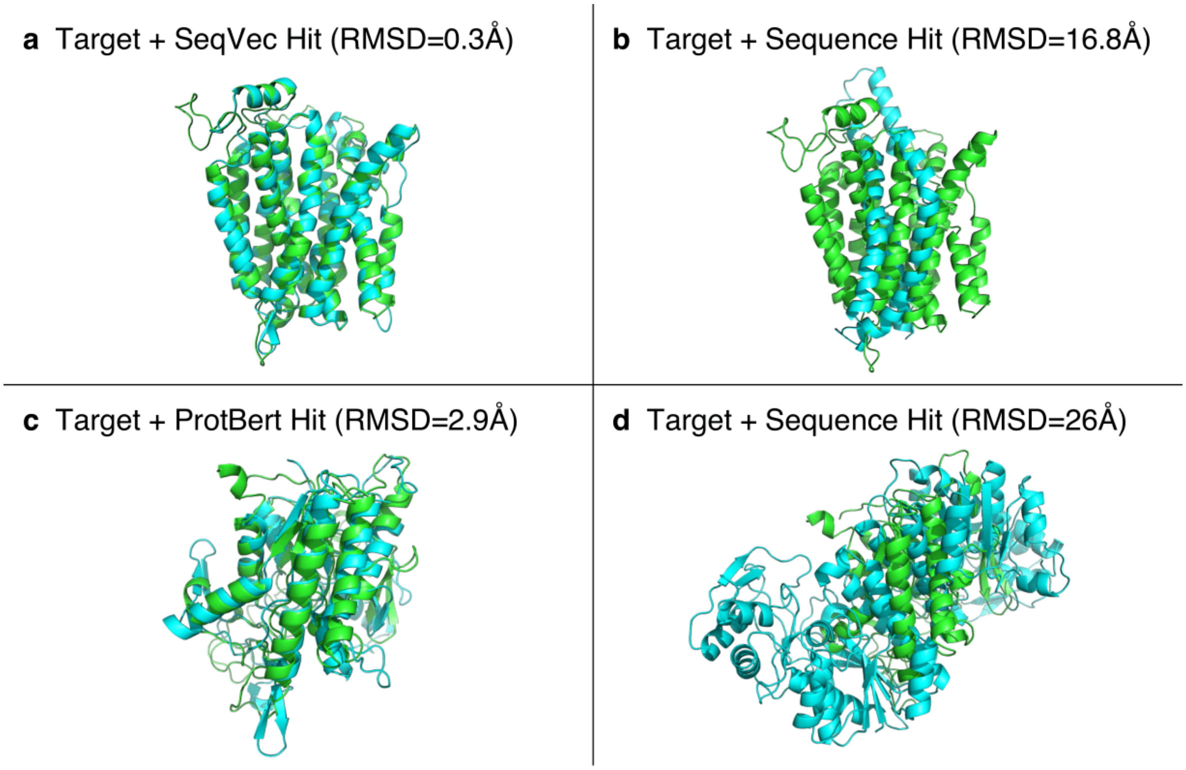

We chose two example targets for which embedding-based inference worked better than homology-based inference and where the embedding similarity was high while the sequence similarity was low. This resulted in two protein triplets (target, embedding-similar hit, sequence-similar hit). For none of the six proteins, structures were available. **a.** Comparative modeling using Swiss-Model[2] mapped the first target with UniProt identifier P40445 and the corresponding hit found using SeqVec embeddings (P53322) to the same structural template of the D-galactonate-proton symporter of *E. coli* (6E9N[3,4]) with the resulting Swiss-Model models having a root-mean-square deviation (RMSD) of 0.3Å. **b.** The sequence-similar hit (Q7PMG3) for the same target mapped to the PDB structure 6C70 (Cryo-EM structure of Orco)[3,5] with RMSD=16.8Å to the structure of the target. **c.** For the second target (Q9STM6), the embedding-based hit (Q9SYF0) was found using ProtBert. Comparative modeling mapped target and hit to PDB structures 3KVN (autotransporter EstA from *Pseudomonas aeruginosa*)[3,6] and 5XTU (GDSL esterase of photobacterium sp. J15)[3,7] with RMSD=2.9Å. **d.** The modeled structure for the corresponding sequence-similar hit (Q564Q1) was based on the PDB structure 1KVQ (UDP-galactose 4-epimerase complexed with UDP-phenol)[3,8] with RMSD=26Å to the structure model of the target.

#### Weak correlation between embedding similarity and GO term similarity

Our method relies on the assumption that proteins with similar embeddings share similar GO annotations. To evaluate this assumption, we randomly chose 5,000 proteins from the *GOA2017-100* set and calculated pairwise distances between these proteins and the remaining proteins in *GOA2017-100* (298,984 proteins) using SeqVec embeddings. We extracted the first 1,000 hits to avoid that our analysis is dominated by random noise for very distant protein pairs. This resulted in 5,000,000 pairs of proteins. For each of those pairs, we calculated the GO term similarity using the Jaccard index (Eqn. 7 in the main text).

The absolute embedding similarity does not necessarily have to correlate with the GO term similarity, i.e. if two proteins have a distance $d_1$, their GO annotations are not necessarily more similar than for two other proteins with a larger distance $d_2$. We rather assume that for a given query, the protein closest to it is more likely to have more similar GO terms than another hit farther away. Also, how similar the annotations of the most similar hit are differs between proteins. There are proteins for which we find hits in the lookup data set with identical annotations while for others, even the most similar hit might still be annotated to very different GO terms. Since both embedding distance and GO term similarity are not directly comparable between proteins, we converted our results to ranks, i.e. for a given query protein, we ordered all hits by distance and gave the hit with the smallest distance rank 1, the hit with the second-smallest distance rank 2, and so forth. We applied the same approach for the GO term similarity. For a perfect correlation, the closest protein (rank 1) should also have the most similar GO annotations (rank 1) and the most distant protein (rank 1000) should have the most dissimilar GO annotations (rank 1000).

For a query protein, its most similar hits (i.e. low ranks) are also more likely to be amongst the closet hits while more dissimilar hits also tend to be further away from the query in embedding space (Fig. S1). This observation corresponds to a weak correlation of $\rho$=0.28 (Spearman's correlation coefficient, p-value < 2.2e-16) between embedding similarity and GO term similarity. This weak correlation was expected to some extent. While we assume that proteins close in embedding space should have similar GO annotations, this does not necessarily imply that proteins far away in embedding space have dissimilar GO annotations.

**Fig. S7: Embedding similarity for different levels of GO term similarity**

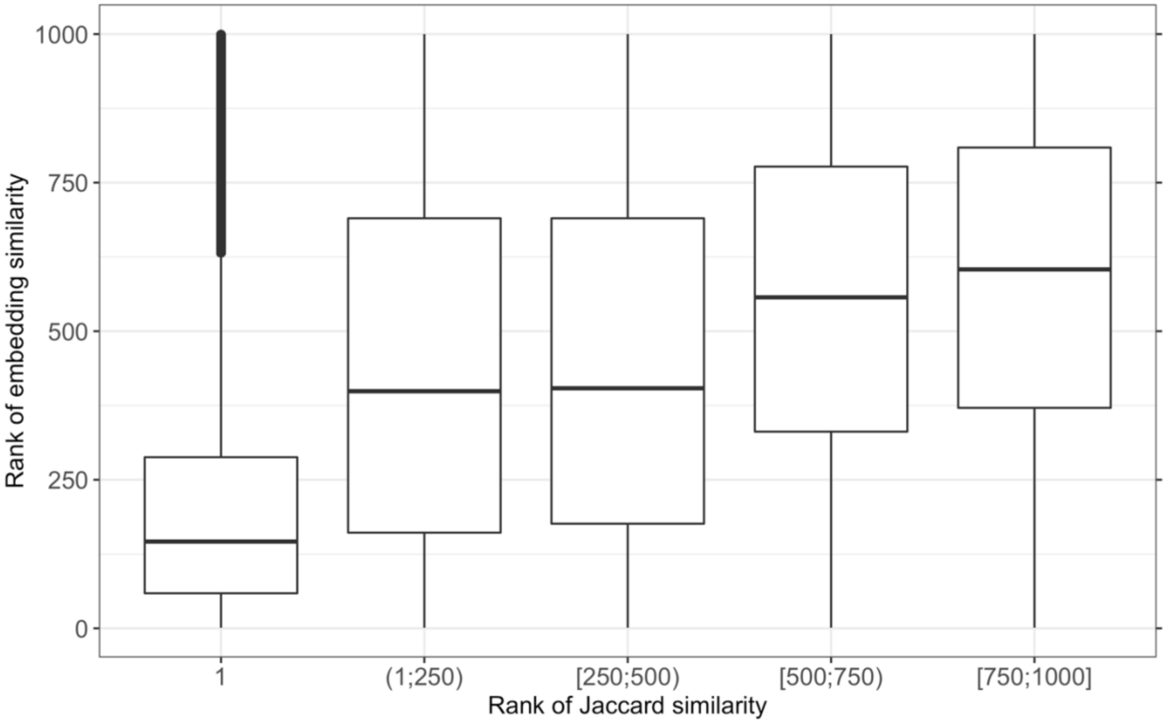

For each query protein, we extracted the 1000 closest proteins and assigned each hit a rank (i) according to its distance to the query (rank 1 = closest hit) and (ii) according to its Jaccard similarity between the GO annotations (rank 1 = most similar annotations). For a set of 5,000 proteins, the ranks for the Jaccard similarity were grouped as shown on the x-axis. On the y-axis, the ranks for the embedding distance are shown. For most proteins, the hit with the most similar annotation (most left box) is also one of the closest hits in embedding space.

**Fig. S8: Fraction of proteomes with predicted GO terms using lookup set *GOA2020***

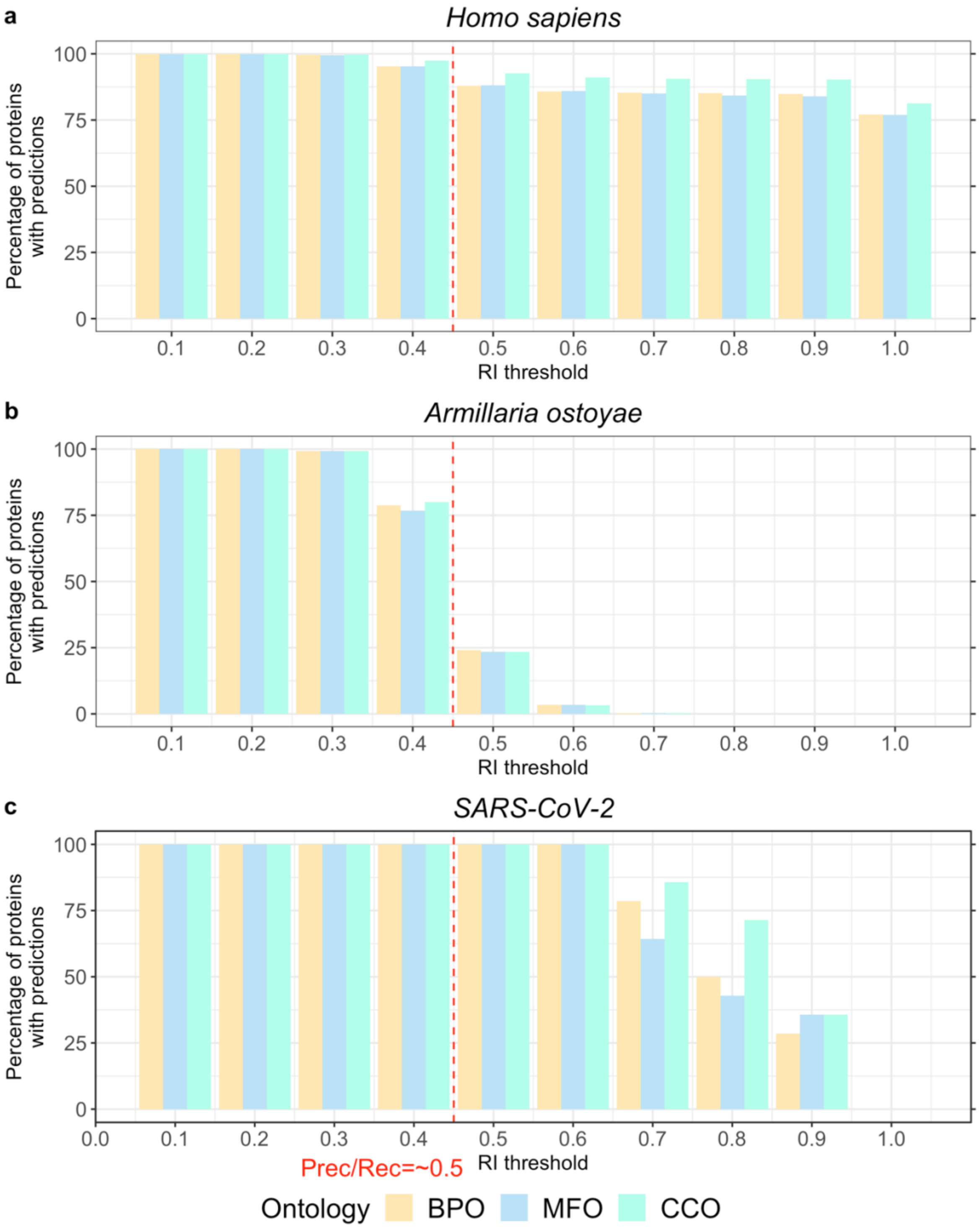

We applied our method to three proteomes (animal: Homo sapiens, fungus: *Armillaria ostoyae*, and virus: SARS-CoV-2) and monitored the fraction of proteins in each proteome for which our method predicted GO terms for different thresholds in embedding similarity (RI, Eqn. 5 in main text). Intervals labelled by number (N=[0.1,1.0]) average over all predictions (with N-0.1<RI≤N). For RI=0.5, our method is expected to achieve roughly 50% precision and recall in all three ontologies (indicated by red dashed line). We used the set GOA2020 as lookup data set which also contains GO annotations not experimentally verified. **a.** The human proteome is well-studied (all 20,370 proteins are in Swiss-Prot) and for many proteins, GO term predictions can be obtained through self-hits, i.e. the annotations are taken from the query protein. Also, for proteins without any GO annotation, our method could predict GO terms at very high reliability probably because there are many well-studied model organisms with experimentally annotated orthologous proteins. **b.** The proteome of the fungus *Armillaria ostoyae* appears more exotic (0.01% of the 22,192 proteins were in Swiss-Prot); high-reliability (RI>0.7) predictions of GO terms were available for few proteins, and for RI>0.5, GO terms were predicted for fewer than half of the proteome. **c.** While annotations were unknown for most proteins of the novel virus SARS-CoV-2 (no coverage at RI=1), many annotations could be transferred from the human SARS coronavirus (SARS-CoV) and the bat coronavirus HKU3 (BtCoV) allowing GO term predictions for all proteins at reliability values as high as RI=0.6.

**Fig. S9: Fraction of proteomes with predicted GO terms using lookup set *GOA2020X***

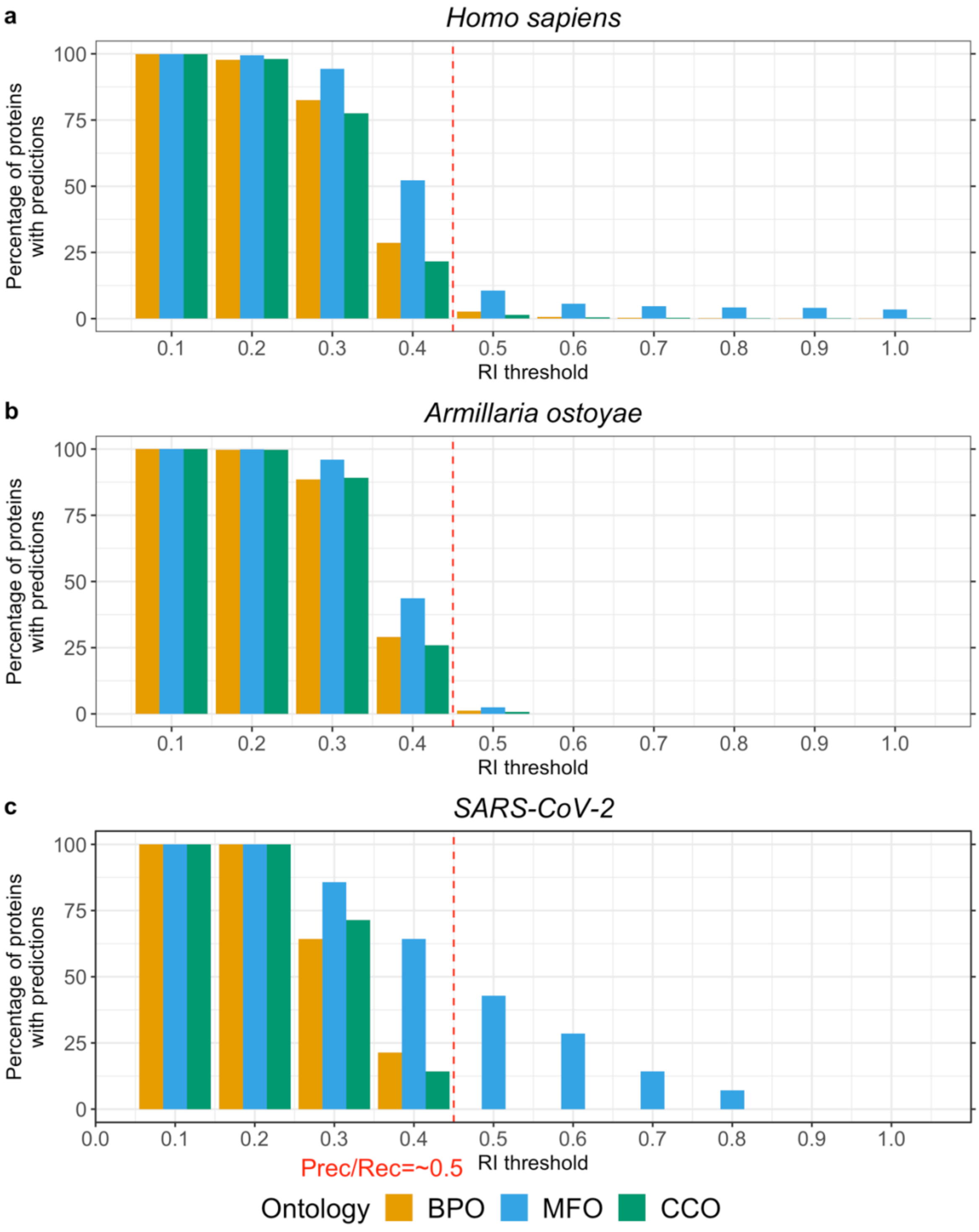

We applied our method to three proteomes (animal: Homo sapiens, fungus: *Armillaria ostoyae*, and virus: SARS-CoV-2) as described in Fig. S6. Instead of *GOA2020*, we used

*GOA2020X* as lookup data set which only contains experimentally verified annotations. **a.** While the human proteome is in general well-studied, most proteins lack GO annotations (almost no annotation transfer at RI=1.0). **b.** For the proteome of the fungus *Armillaria ostoyae*, almost no experimental GO annotations could be transferred at high reliability (RI>0.5). **c.** While SARS-CoV-2 is a very novel and therefore not well-studied proteome, for almost 50% of the proteins, GO annotations for MFO could be inferred at RI>0.5 from the human SARS coronavirus (SARS-CoV) and the bat coronavirus HKU3 (BtCoV) which are both more well-studied.

#### Fig. S10: Visualization of predicted GO term for Nsp7b from SARS-CoV-2

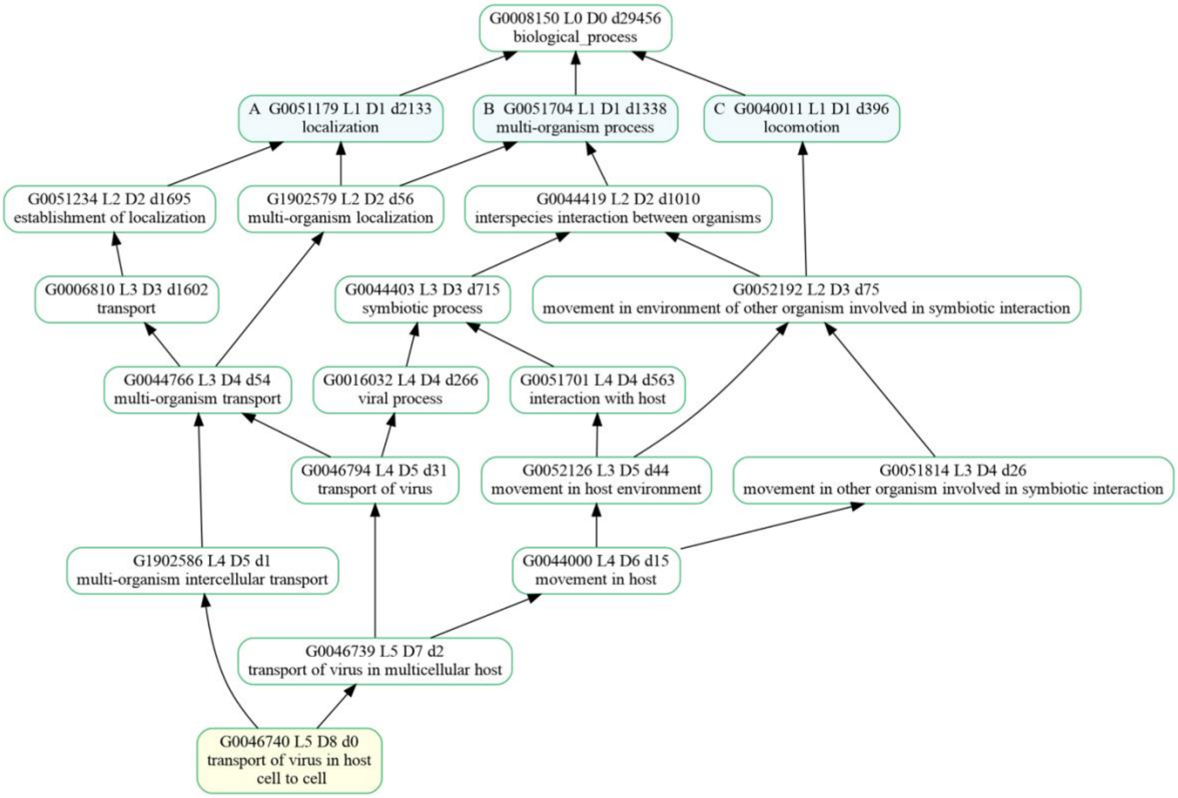

To further analyze the predicted GO terms, the leaf terms can be visualized in the GO hierarchy. For the prediction of “GO:0046740” for the non-structural protein 7b from SARS-CoV-2, this visualization revealed that the functionality of this protein constituted two main components: The interaction with the host and the actual transportation.

**Fig. S11: Visualization of the embedding generation using SeqVec.**

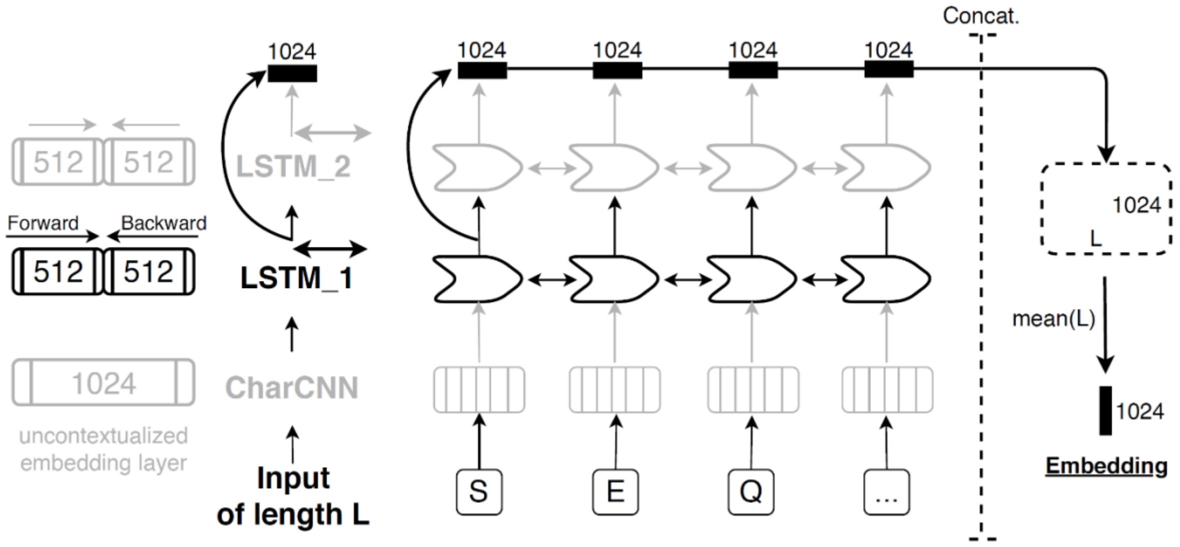

We outline the process of generating fixed-size embeddings for protein sequences of variable length using SeqVec. The example illustrated here shows how the first three residues of protein sequence ("SEQ..") are processed: first, the three layers of SeqVec (uncontextualized: CharCNN; contextualized: LSTM layer 1 and LSTM layer 2) project the protein sequence to vector space. The CharCNN generates vectors of size 1024 without considering neighboring residues. In contrast to this, the two LSTM layers process the sequence in both directions, each creating a vector of size 512. The vectors of both directions are concatenated for each LSTM independently, resulting in an embedding size of 1024 for each LSTM layers. In a second step, only embeddings of the first LSTM layer are extracted, concatenated and averaged over the length of the protein (global average pooling), resulting in a 1024-dimensional embedding.

**Table S1: Correlation between $F_{max}$ and protein length**

| | Spearman's correlation coefficient | P-value |
|---|---|---|
| **SeqVec** | -0.03 | 0.27 |
| **ProtBert** | -0.03 | 0.16 |
| **BLAST** | 0.05 | 0.06 |

* Global average pooling for long sequences could lead to information loss because important information might be averaged out if the signal is not consistent over the protein length. However, we did not observe a correlation between protein length and performance for either SeqVec or ProtBert. The same holds true for homology-based inference ("BLAST"). Correlation was measured using Spearman's rank correlation coefficient.

**Table S2: $F_{max}$ and average number of predicted GO terms for different values of *k***

| | $F_{max}$ | | |
|---|---|---|---|
| | **BPO** | **MFO** | **CCO** |
| **k=1** | 37±2% | 50±3% | 57±2% |
| **k=2** | 37±2% | 51±2% | 58±2% |
| **k=3** | 37±2% | 50±2% | 58±2% |
| **k=4** | 37±2% | 51±2% | 58±2% |
| **k=5** | 36±2% | 51±2% | 59±2% |
| **k=10** | 36±2% | 49±3% | 56±2% |

* The number of neighbors included for the annotation transfer did not affect $F_{max}$. Evaluation was performed for the no-knowledge (NK) set of proteins for which no annotations in any ontology were available at the submission deadline of CAFA3. Error estimates indicate 95% confidence intervals.

**Table S3: Precision, recall, and average number of predicted GO terms for different values of *k***

| | Precision (shown as percentages) | | | Recall (shown as percentages) | | | Average number of predicted GO terms per protein | | |
|---|---|---|---|---|---|---|---|---|---|
| | **BPO** | **MFO** | **CCO** | **BPO** | **MFO** | **CCO** | **BPO** | **MFO** | **CCO** |
| **k=1** | 32±2 | 47±3 | 54±3 | 43±3 | 54±3 | 62±3 | 46.5 | 11.7 | 15.7 |
| **k=2** | 28±2 | 40±3 | 49±3 | 49±2 | 60±3 | 69±3 | 66.3 | 15.4 | 19.8 |
| **k=3** | 25±2 | 37±3 | 46±2 | 52±2 | 62±3 | 72±3 | 80.6 | 18.0 | 22.7 |
| **k=4** | 23±2 | 35±3 | 43±2 | 55±2 | 64±3 | 75±2 | 97.6 | 20.1 | 25.8 |
| **k=5** | 21±2 | 32±3 | 40±2 | 57±2 | 66±3 | 78±2 | 113.5 | 22.4 | 29.0 |
| **k=10** | 16±1 | 24±2 | 30±2 | 65±2 | 72±3 | 83±2 | 183.5 | 33.4 | 43.2 |

* The average number of predicted GO terms per protein increased with increasing values of *k* when all predictions are taken into account. This increase in the number of predicted terms led to a decrease in precision, but to an increase in recall. Taking more proteins than the closest one into account can be beneficial to e.g., increase the specificity and quality of the predicted terms if only a few, unspecific terms are annotated to the closest hit. Evaluation was performed for the no-knowledge (NK) set of proteins for which no annotations in any ontology were available at the submission deadline of CAFA3. Error estimates indicate 95% confidence intervals.

**Table S4: Fmax for different combinations of embedding-based and homology-based annotation transfer**

| | $F_{max}$ | | |
|---|---|---|---|
| | BPO | MFO | CCO |
| SeqVec-2020 | 51±2% | 61±3% | 65±2% |
| SeqVec/ProtBert | 51±2% | 60±2% | 66±2% |
| SeqVec/BLAST | 50±2% | 61±2% | 65±2% |
| ProtBert/BLAST | 49±2% | 60±2% | 65±2% |
| SeqVec/ProtBert/BLAST | 51±2% | 61±2% | 66±2% |

* Combining embedding-based annotation transfer for different language models (SeqVec, ProtBert) and homology-based transfer (BLAST) did not improve performance over embedding-based transfer using SeqVec (SeqVec-2020). For this combination, every term predicted for either of the methods was also include in the final, combined prediction. Maybe this approach is too simple to reflect the complex relationship between embedding similarity, sequence similarity and prediction quality and only a more sophisticated combination could improve over the single method (SeqVec).

#### Table S5: Annotated and predicted GO terms for SARS-CoV-2 proteins

| | Annotated | | | Predicted | | |
|---|---|---|---|---|---|---|
| | **BPO** | **MFO** | **CCO** | **BPO** | **MFO** | **CCO** |
| **P0DTC1** | GO:0006508<br>GO:0016032<br>GO:0030683<br>GO:0039502<br>GO:0039503<br>GO:0039520<br>GO:0039548<br>GO:0039579<br>GO:0039595<br>GO:0039648<br>GO:0039657<br>GO:0090305 | GO:0003723<br>GO:0004518<br>GO:0004519<br>GO:0008233<br>GO:0008234<br>GO:0016787<br>GO:0036459<br>GO:0046872 | GO:0016020<br>GO:0016021<br>GO:0030430<br>GO:0033644<br>GO:0044220 | GO:0001172<br>GO:0006508<br>GO:0019079<br>GO:0019082<br>GO:0039502<br>GO:0039520<br>GO:0039548<br>GO:0039579<br>GO:0039595<br>GO:0039648<br>GO:0090305 | GO:0036459<br>GO:0003723<br>GO:0003968<br>GO:0004197<br>GO:0004519<br>GO:0008242<br>GO:0008270 | GO:0016020<br>GO:0016021<br>GO:0030430<br>GO:0033644<br>GO:0044220 |
| **P0DTD1** | GO:0001172<br>GO:0006508<br>GO:0016032<br>GO:0030683<br>GO:0032259<br>GO:0032508<br>GO:0039502<br>GO:0039503<br>GO:0039520<br>GO:0039579<br>GO:0039595<br>GO:0039644<br>GO:0039648<br>GO:0039657<br>GO:0090305 | GO:0000166<br>GO:0003678<br>GO:0003723<br>GO:0003724<br>GO:0003968<br>GO:0004386<br>GO:0004518<br>GO:0004519<br>GO:0004527<br>GO:0005524<br>GO:0008168<br>GO:0008233<br>GO:0008234<br>GO:0016740<br>GO:0016779<br>GO:0016787<br>GO:0036459<br>GO:0046872 | GO:0016020<br>GO:0016021<br>GO:0030430<br>GO:0033644<br>GO:0044172<br>GO:0044220 | GO:0001172<br>GO:0006351<br>GO:0006508<br>GO:0019082<br>GO:0019083<br>GO:0032259<br>GO:0032508<br>GO:0039502<br>GO:0090503<br>GO:0039520<br>GO:0039579<br>GO:0039595<br>GO:0039644<br>GO:0039648<br>GO:0039694 | GO:0000175<br>GO:0003678<br>GO:0003723<br>GO:0003724<br>GO:0003968<br>GO:0004197<br>GO:0004519<br>GO:0005524<br>GO:0008168<br>GO:0008242<br>GO:0008270<br>GO:0036459<br>GO:0042802 | GO:0016020<br>GO:0016021<br>GO:0030430<br>GO:0033644<br>GO:0044172<br>GO:0044220 |
| **P0DTC2** | GO:0009405<br>GO:0016032<br>GO:0019062<br>GO:0039654<br>GO:0039663<br>GO:0044650<br>**GO:0046718** | **GO:0005515**<br>**GO:0046789** | GO:0016020<br>GO:0016021<br>GO:0019012<br>GO:0019031<br>GO:0020002<br>GO:0033644<br>GO:0044173<br>GO:0055036 | GO:0009405<br>GO:0019064<br>GO:0039654<br>**GO:0046718**<br>GO:0046813<br>GO:0075509 | GO:0042802<br>**GO:0046789** | GO:0016021<br>GO:0019012<br>GO:0019031<br>GO:0020002<br>GO:0044173<br>GO:0055036 |
| **P0DTC3** | No annotations | | GO:0005576<br>GO:0016020<br>GO:0016021<br>GO:0019012<br>GO:0020002<br>GO:0030430<br>GO:0033644<br>GO:0044177<br>GO:0044178 | GO:0034220<br>GO:0039707<br>GO:0051259 | GO:0005216 | GO:0005576<br>GO:0016020<br>GO:0016021<br>GO:0019012<br>GO:0020002<br>GO:0030430<br>GO:0044177<br>GO:0044178<br>GO:0044385 |

| | Annotated | | | Predicted | | |
|---|---|---|---|---|---|---|
| | **BPO** | **MFO** | **CCO** | **BPO** | **MFO** | **CCO** |
| **P0DTC4** | No annotations | | GO:0016020<br>GO:0016021<br>GO:0033644<br>GO:0044177<br>GO:0044178 | GO:0044662<br>GO:0046760 | GO:0015078 | GO:0016020<br>GO:0016021<br>GO:0044172<br>GO:0044177<br>GO:0044178 |
| **P0DTC5** | GO:0016032<br>GO:0030683 | GO:0039660 | GO:0016020<br>GO:0016021<br>GO:0019012<br>GO:0019031<br>GO:0033644<br>GO:0044177<br>GO:0044178<br>GO:0055036 | GO:0019058<br>GO:0030683 | GO:0039660 | GO:0016021<br>GO:0019012<br>GO:0019031<br>GO:0030430<br>GO:0044177<br>GO:0044178<br>GO:0055036 |
| **P0DTC6** | GO:0009405 | No annotations | GO:0016020<br>GO:0033644<br>GO:0044165<br>GO:0044167<br>GO:0044177<br>GO:0044178 | GO:0009405 | GO:0005125<br>GO:0005126 | GO:0016020<br>GO:0044165<br>GO:0044167<br>GO:0044177<br>GO:0044178 |
| **P0DTC7** | GO:0016032<br>GO:0039646<br>GO:0060153 | No annotations | GO:0016020<br>GO:0016021<br>GO:0019012<br>GO:0033644<br>GO:0044165<br>GO:0044167<br>GO:0044173<br>GO:0044177<br>GO:0044178 | GO:0039646 | GO:0005515<br>GO:0005537 | GO:0016020<br>GO:0016021<br>GO:0019012<br>GO:0044165<br>GO:0044167<br>GO:0044173<br>GO:0044177<br>GO:0044178 |
| **P0DTD8** | No annotations | | GO:0016020<br>GO:0016021<br>GO:0033644 | GO:0046740 | GO:0015078 | GO:0016020<br>GO:0016021<br>GO:0033644 |
| **P0DTC8** | No annotations | | | GO:0006954<br>GO:0031666<br>GO:0045087 | GO:0005515 | GO:0005576<br>GO:0005615 |
| **P0DTC9** | No annotations | GO:0003723 | GO:0019012<br>GO:0019013<br>GO:0044172<br>GO:0044177 | GO:0000413<br>GO:0006457<br>GO:0016567 | GO:0003723 | GO:0019012<br>GO:0019013<br>GO:0030430<br>GO:0044172<br>GO:0044177<br>GO:0044220 |
| **P0DTD2** | No annotations | | GO:0016020<br>GO:0030430<br>GO:0033644<br>GO:0044161<br>GO:0044162 | GO:0006412 | GO:0000049<br>GO:0003735 | GO:0016020<br>GO:0030430<br>GO:0044161<br>GO:0044162 |
| **A0A663DJA2** | No annotations | | | GO:0009734<br>GO:0010930<br>GO:0048364 | GO:0005506<br>GO:0009055<br>GO:0020037 | GO:0016020<br>GO:0016021 |

* Of the 14 proteins of SARS-CoV in UniProt, 7 have annotations in BPO, 6 in MFO and 12 in CCO. Of the 138 GO terms annotated, only three are experimentally verified. We predicted 83% of the annotations for BPO, 47% for MFO, and 87% for CCO. Bold terms indicate experimentally verified annotations; underlined terms indicate predicted terms which are also annotated.

**Table S6: Datasets for similarity lookup at different sequence identity thresholds.**

| | GOA2017 | GOA2020 | %identical proteins | Average sequence identity |
|---|---|---|---|---|
| **Full** | 307,287 | 295,558 | | |
| **100% Seq. ID.** | 303,984 | 292,059 | 87% | 44% |
| **90% Seq. ID.** | 302,052 | 290,030 | 87% | 44% |
| **80% Seq. ID.** | 300,571 | 288,561 | 87% | 44% |
| **70% Seq. ID.** | 298,677 | 286,748 | 87% | 44% |
| **60% Seq. ID.** | 295,823 | 284,043 | 87% | 43% |
| **50% Seq. ID.** | 290,094 | 278,711 | 87% | 43% |
| **40% Seq. ID.** | 281,539 | 270,691 | 87% | 43% |
| **30% Seq. ID.** | 262,440 | 260,056 | 72% | 43% |
| **20% Seq. ID.** | 242,154 | 242,261 | 63% | 43% |

* The first two columns show the size of data sets extracted from the Gene Ontology Annotation (GOA) database [9-11] in January 2017 (GOA2017) and January 2020 (GOA2020). The first row shows the full data set when only considering sequences from Swiss-Prot [12] and removing proteins only annotated to the roots of the three ontologies. The remaining rows represent data sets redundancy reduced against the CAFA3 targets at sequence identity thresholds of 100, 90, 80, 70, 60, 50, 40, 30 and 20%, respectively. Redundancy reduction was performed using CD-HIT and PSI-CD-HIT [13,14]. The last two columns show the agreement between the two GOA versions, *%identical proteins* gives the percentage of identical proteins in both sets (by UniProt identifier), and *Average sequence identity* states the average sequence identity of all protein pairs.

### References for Supporting Online Material

1 Zhou, N. *et al.* The CAFA challenge reports improved protein function prediction and new functional annotations for hundreds of genes through experimental screens. *Genome Biol* **20**, 244, doi:10.1186/s13059-019-1835-8 (2019).

2 Waterhouse, A. *et al.* SWISS-MODEL: homology modelling of protein structures and complexes. *Nucleic Acids Res* **46**, W296-W303, doi:10.1093/nar/gky427 (2018).

3 Berman, H. M. *et al.* The Protein Data Bank. *Nucleic Acids Res* **28**, 235-242, doi:10.1093/nar/28.1.235 (2000).

4 Leano, J. B. *et al.* Structures suggest a mechanism for energy coupling by a family of organic anion transporters. *PLoS Biol* **17**, e3000260, doi:10.1371/journal.pbio.3000260 (2019).

5 Butterwick, J. A. *et al.* Cryo-EM structure of the insect olfactory receptor Orco. *Nature* **560**, 447-452, doi:10.1038/s41586-018-0420-8 (2018).

6 van den Berg, B. Crystal structure of a full-length autotransporter. *J Mol Biol* **396**, 627-633, doi:10.1016/j.jmb.2009.12.061 (2010).

7 Mazlan, S. *et al.* Crystallization and structure elucidation of GDSL esterase of Photobacterium sp. J15. *Int J Biol Macromol* **119**, 1188-1194, doi:10.1016/j.ijbiomac.2018.08.022 (2018).

8 Thoden, J. B., Gulick, A. M. & Holden, H. M. Molecular structures of the S124A, S124T, and S124V site-directed mutants of UDP-galactose 4-epimerase from Escherichia coli. *Biochemistry* **36**, 10685-10695, doi:10.1021/bi9704313 (1997).

9 *GOA*, <http://www.ebi.ac.uk/GOA> (2020).

10 Camon, E. *et al.* The Gene Ontology Annotation (GOA) Database: sharing knowledge in Uniprot with Gene Ontology. *Nucleic Acids Res* **32**, D262-266, doi:10.1093/nar/gkh021 (2004).

11 Huntley, R. P. *et al.* The GOA database: gene Ontology annotation updates for 2015. *Nucleic Acids Res* **43**, D1057-1063, doi:10.1093/nar/gku1113 (2015).

12 Boutet, E., Lieberherr, D., Tognolli, M., Schneider, M. & Bairoch, A. UniProtKB/Swiss-Prot. *Methods Mol Biol* **406**, 89-112, doi:10.1007/978-1-59745-535-0_4 (2007).

13 Fu, L., Niu, B., Zhu, Z., Wu, S. & Li, W. CD-HIT: accelerated for clustering the next-generation sequencing data. *Bioinformatics* **28**, 3150-3152, doi:10.1093/bioinformatics/bts565 (2012).

14 Li, W. & Godzik, A. Cd-hit: a fast program for clustering and comparing large sets of protein or nucleotide sequences. *Bioinformatics* **22**, 1658-1659, doi:10.1093/bioinformatics/btl158 (2006).
